# Detecting Nuclear Pore Complex assembly in living cells

**DOI:** 10.1101/2025.07.30.667432

**Authors:** Annemiek C Veldsink, Jonas S Fischer, Hanna M Terpstra, Philip J Mannino, Eline MF de Lange, Sophie Hell, Koen J van Benthem, Leila J Saba, Anton Steen, Rifka Vlijm, Matthias Heinemann, Karsten Weis, C Patrick Lusk, Liesbeth M Veenhoff

## Abstract

The formation of nuclear pore complexes (NPCs) — vital gateways regulating nuclear-cytoplasmic transport — is a highly orchestrated process requiring the integration of hundreds of nucleoporins into the nuclear envelope. A major challenge in studying this assembly process in living cells has been the difficulty to distinguish newly forming NPCs from their mature counterparts. Here, we present a powerful nanobody-based approach that overcomes this limitation. We demonstrate that a nanobody targeting the nucleoporin Nic96 from *Saccharomyces cerevisiae* selectively binds newly synthesized Nic96 subcomplexes prior to its incorporation into NPCs *in vivo*. Importantly, nanobody-bound Nic96 is incorporated in NPCs, and expression of the nanobody does not disrupt nuclear transport, cell growth, or lifespan, nor does it show genetic interactions with known NPC assembly surveillance pathways — making it an ideal, non-perturbing tool to study NPC biogenesis in yeast. Illustrating the use of the Nic96 nanobody we report novel aspects of the early stages of assembly, including co-recruitment of Kap121 and VHH[Nic96] to putative assembly sites. In addition, we show local enrichment of newly synthesized nucleoporins on nuclear envelope proximal lipid droplets, including a specific subset of Pdr16- and Ldo16-positive lipid droplets near the nucleus-vacuole junction. When NPC assembly is delayed, association of Nic96 subcomplexes with lipid droplets increases. These findings illustrate how the strategy to pulse-label newly forming NPCs opens new avenues for dissecting the spatiotemporal regulation of NPC assembly and misassembly.

## Introduction

Nuclear pore complexes (NPCs) are large and broadly conserved structures embedded in the nuclear envelope (NE) of all eukaryotes ^1^. NPCs allow macromolecular trafficking between the nucleus and cytosol ^2,3^ and function as organizational hubs at the nuclear periphery, docking for example DNA lesions and the proteasome ^4^. The importance of proper NPC function and/or adequate NPC numbers is underscored by recent research in disease contexts. For instance, some cancer cells have higher NPC numbers and show increased sensitivity to interventions that disrupt NPC assembly ^5^. Vice versa, in neurodegenerative diseases where NPC function is compromised, increasing NPC number alleviates several disease-related phenotypes ^6^.

NPCs assemble into a highly conserved, cylindrical structure that is built from hundreds of proteins called nucleoporins (Nups). The NPC’s scaffold is organized in eight rotationally symmetrical spokes, that connect to the NE membrane and the NPC’s membrane ring. In the plane of the membrane, the NPC is composed of inner and outer ring structures. The outer rings on the cytoplasmic and nuclear sides are formed by the Nups from the Y-complexes, amongst which *Saccharomyces cerevisiae* Nup84 (hNup107). Flexible linker Nups tie together the spokes and ring structures ^7,8^, thereby allowing for flexibility in the structure that can accommodate dilation and contraction ^9,10^. Intrinsically disordered Nups that are enriched in repeats of phenylalanine and glycine, the FG-Nups, form the permeability barrier. The Nsp1-Nup57-Nup49 subcomplex, the central transport Nup (CTN) trimer, is anchored to the inner rings through Nic96 (hNup93) ^11,12^. Asymmetry in the NPC structure is introduced by Nups from the cytoplasmic complex that assemble on the cytosolic side and the nuclear basket Nups that dock on the nuclear side.

In dividing budding yeast cells, approximately one hundred NPCs assemble during each cell cycle to ensure that the daughter cells receive their share of NPCs ^13^. Given the essential nature of NPCs, any perturbations in this process typically allow only a few more divisions before the cells die ^14,15^. Assembling a new NPC is an immense challenge: many Nups contain repetitive structural modules and disordered regions that need to be incorporated in correct stoichiometries at the NE, all while simultaneously remodelling the NE to create the pore. To orchestrate the assembly process, many subcomplexes assemble co-translationally ^16,17^, preventing aberrant interactions. Some Nups even translate locally at the NPC ^16^. In mammalian cells, the assembly of NPCs also relies on the molecular chaperone DNAJB6 that delays the transition to aggregated forms of the FG-Nups ^18,19^. These mechanisms provide a glimpse of the complexity of the process, but the molecular events that initiate subcomplex assembly as well as the subsequent steps leading to the assembly of the higher-order, NE-embedded structure remain poorly defined.

A major difficulty that hampers a better understanding of the mechanisms guarding the (pre-)assembly process is the absence of tools to discriminate between assembly intermediates and mature NPCs. Genomically encoded fluorophore switches (e.g. RITE cassette ^20,21^) are amenable to live-cell imaging and can assess NPC maturation over timescales of hours or days, but they cannot capture early assembly events. Current approaches capable of detecting early assembly events rely on biochemical assays (KARMA ^22,23^), or ex vivo assembly strategies ^24^, but tools to capture NPC assembly dynamics in living cells are scarce ^25,26^ and absent for systems probing the assembly of endogenous Nups. Such tools require approaches for NPC labelling that allow characterization of nascent and mature NPC populations in the NE of living cells. In this study, we show that a previously developed nanobody against budding yeast Nic96 ^27^, whose epitope is shielded in mature NPCs, can report on NPC assembly events in living yeast cells by preferentially binding accessible Nic96 during pre-assembly and assembly stages rather than binding Nic96 in mature NPCs. Our biochemical and live-cell imaging data demonstrates that VHH[Nic96] integrates into Nic96-containing subcomplexes and can be used to monitor NPC assembly without impairing NPC function. Using this tool, we observed co-recruitment of Kap121 and VHH[Nic96] to putative assembly sites and we identified newly synthesized Nic96 subcomplexes near lipid droplets at the NE, with this spatial association strongly dependent on assembly efficiency. Together, these findings establish VHH[Nic96] as a valuable probe for spatiotemporal analysis of NPC assembly in living yeast cells.

## Results

### VHH[Nic96] binds newly synthesized Nic96 and subcomplex members

To detect NPC assembly *in vivo, w*e reasoned that a nanobody (single-domain antibody) targeting a Nup epitope that is inaccessible in the mature NPC structure could serve as a valuable tool. In this case, nanobody binding would conceivably be biased towards the pre-assembly and assembly stage where the epitope is accessible. The VHH-SAN12 nanobody that was previously developed for structural biology recognizes the C-terminal domain of the inner ring component Nic96 (from here on VHH[Nic96]) with high specificity and nanomolar affinity *in vitro* ^27^. According to recent structures of the yeast NPC ^28^, the epitope of VHH[Nic96] is buried deep within the NPC structure (Fig. 1A) and thus conceivably shielded, therefore making VHH[Nic96] a potential candidate for labelling NPC assembly events (Fig. 1B). As a reference for an NPC-targeting nanobody that does not show specificity towards the (pre)assembly stage, we used VHH-SAN8 ^27^, which recognizes an accessible epitope on the outer ring component Nup84 (from here on VHH[Nup84], Fig. 1A). We previously showed that VHH[Nup84] rapidly labels existing NPCs ^29^ and therefore VHH[Nup84] does not specifically label the assembly stage.

**Fig. 1.**
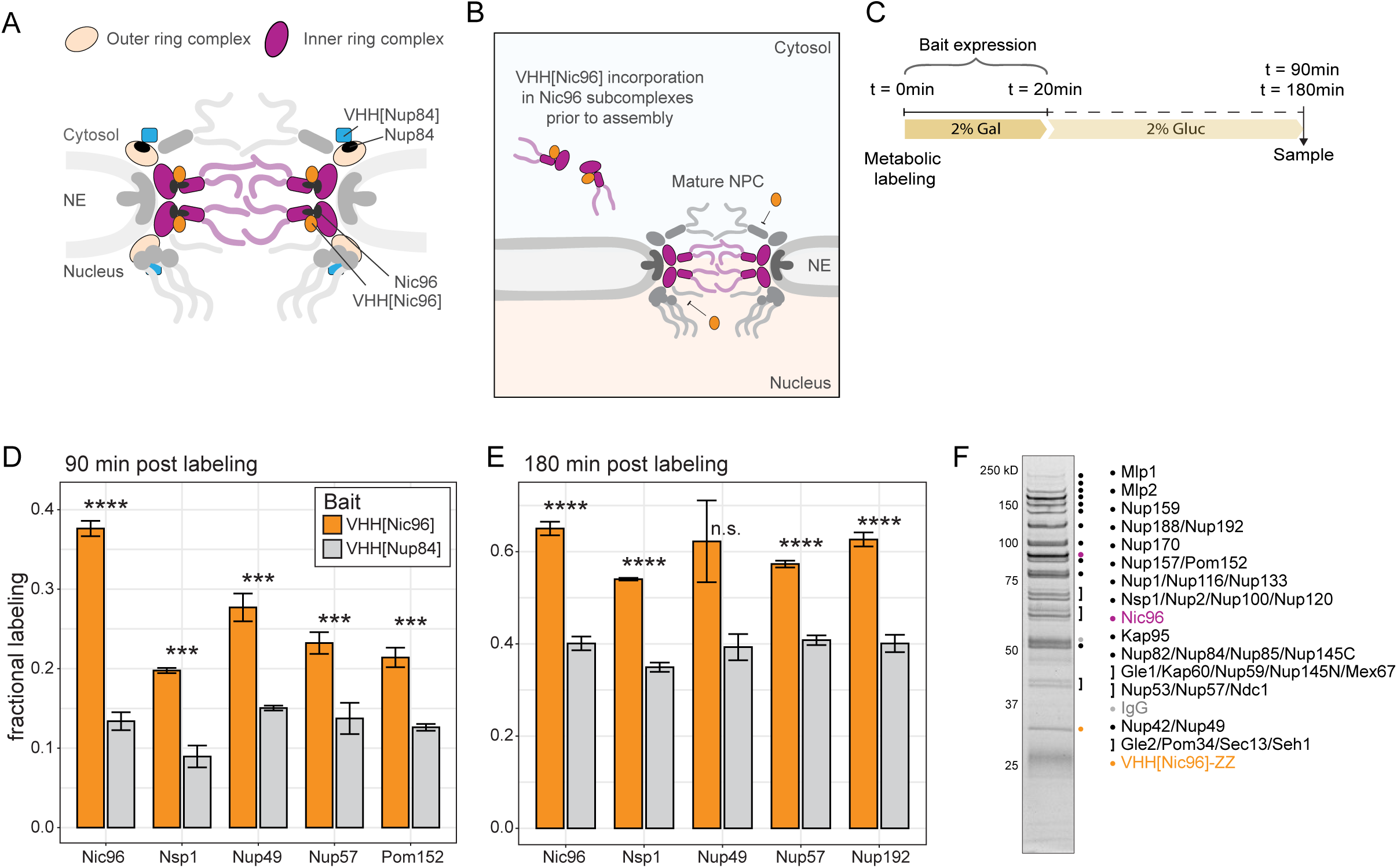
VHH[Nic96] binds newly synthesized Nic96 and direct Nic96 interaction partners **A.** Cartoon of the yeast nuclear pore complex and nanobodies (VHH[Nup84], blue square; VHH[Nic96], orange ellipse) bound to Nup84 and Nic96 (black). The NPC’s outer and inner ring complexes are highlighted. **B.** Cartoon illustrating the hypothesized incorporation of VHH[Nic96] into Nic96 subcomplexes prior to assembly but not mature NPCs due to the shielded Nic96 epitope in mature NPCs. **C.** Experimental set-up in panels D, E. ZZ-tagged VHH[Nic96] and VHH[Nup84] are expressed from an inducible galactose promoter. A short, 20-minute expression burst (addition of 2% galactose (Gal)) was followed by addition of 2% glucose (Gluc) to stop expression and to chase nanobody incorporation in NPCs over time. Nanobody induction was concomitant with metabolic isotope labelling. Affinity purifications were performed at indicated timepoints (t) from the start of the experiment, in two separate experiments. **D.** Fractional labelling (Heavy / (Heavy + Light)) of Nic96, Nsp1, Nup49, Nup57, and Pom152 co-purifying with VHH[Nic96] compared to VHH[Nup84] at t=90min. The median fractional labelling of three biological replicates (bar) and the standard deviation (error bar) are plotted. Significance was assessed using a two-tailed Student’s t-test. See Fig. S1A for all copurified proteins in VHH[Nic96]-ZZ affinity purifications. **E.** As in D but at t=180min showing fractional labelling of Nic96, Nsp1, Nup49, Nup57 and Nup192 co-purifying with VHH[Nic96] compared to VHH[Nup84]. See Fig. S1B for all copurified proteins in VHH[Nic96]-ZZ affinity purifications. **F.** All Nups co-purify in VHH[Nic96]-ZZ affinity purifications (see Fig. S1D). VHH[Nic96] is annotated in orange, Nic96 in purple.

To see if VHH[Nic96] binds new Nic96 and interaction partners of Nic96, we employed the recently developed KARMA (*K*inetic *A*nalysis of incorporation *R*ates in *M*acromolecular *A*ssemblies) approach^22^. We pulsed the expression of VHH[Nic96]-ZZ for 20 minutes whilst simultaneously switching growing yeast cultures from medium containing light lysine to heavy lysine (Fig. 1C). We determined the interactome of VHH[Nic96]-ZZ after an additional 70 or 160 minutes by performing affinity purifications (timepoints 90- or 180-minutes, respectively). This pulse-chase experiment showed that at both timepoints Nic96 and the copurifying CTN trimer (Nup57, Nup49 and Nsp1) had incorporated the heavy lysine (Fig. 1DE). Importantly, the Nic96 and the CTN trimer had a significantly higher fractional labelling (heavy lysine / total lysine) in the VHH[Nic96]-ZZ affinity purifications compared to the VHH[Nup84]-ZZ affinity purifications (Fig. 1DE, see Fig. S1AB for all co-purified proteins). This shows that VHH[Nic96]-ZZ primarily interacts with newly synthesized variants of these Nups. Of note, the pronounced Nup overlabelling in VHH[Nic96] purifications likely represents an underestimation, as intracellular pools of light lysine and light-labelled Nup precursors dampen the fractional labelling ^22^. This data is in line with reports showing that Nic96 interacts with the CTN-subcomplex in a co-translational manner ^17^, and indicates that 1) binding of VHH[Nic96] does not prevent newly synthesized Nic96 from incorporating into this subcomplex and 2) that VHH[Nic96] likely becomes part of this subcomplex prior to its assembly into the full NPC structure. We also detected the transmembrane Nup Pom152, and the inner ring Nup Nup192 – a Nic96 interaction partner in the inner ring ^12,30^ that is part of the same assembly group as Nic96 and the CTN-subcomplex ^22^ – at 90- and 180-minutes post labelling, respectively. Both were also significantly more heavy-labelled than in the control VHH[Nup84]-ZZ affinity purifications (Fig. 1DE). Together, these findings highlight that VHH[Nic96] indeed does not bind fully assembled NPCs but specifically labels newly synthesized Nic96 subcomplexes.

### VHH[Nic96]-mNG is incorporated in NPCs during assembly

In our KARMA experiments, we predominantly detected the direct interaction partners of Nic96.To better understand whether VHH[Nic96] is ultimately incorporated into fully assembled NPCs, after its initial rapid binding to Nic96 and direct binding partners of Nic96 (Fig. 1D), we screened for buffer conditions ^31^ that could maintain the complete VHH[Nic96]-ZZ interactome during affinity purifications (Fig. S1C). Indeed, all Nups from the NPC specifically co-purified with constitutively expressed VHH[Nic96]-ZZ as a bait (Fig. 1F). Vice versa, affinity purifications using endogenously tagged Pom34-ZZ as bait also specifically co-purified VHH[Nic96] with all Nups (Fig. S1E).

To visually examine VHH[Nic96]’s incorporation in NPCs, we fused VHH[Nic96] to mNeongreen (mNG), a bright fluorescent protein with a short maturation time in yeast ^32^ to allow for pulse-chase labelling of Nic96 subcomplexes (Fig. 2A). We first examined VHH[Nic96] localization after a long, overnight expression pulse (Fig. 2B) and under constitutive, Nup expression levels (Fig. S2A). In both cases, VHH[Nic96]-mNG localized to the NE in an NPC-like pattern, although likely sub-stoichiometrically to Nic96 as its total fluorescence was lower than endogenously tagged Nic96-mNG (Fig. 2B). Following a shorter, 20-minute induction pulse, the VHH[Nic96] labelling pattern overlapped with the signal of Nup60-mTQ2 (Fig. 2C). Furthermore, in Nup133ΔN and nup84Δ genetic backgrounds where NPCs cluster in the NE ^33,34^, the VHH[Nic96]-mNG signal also clustered and overlapped with NPCs in the NE (Fig. 2D, Fig. S2BC).

**Fig. 2.**
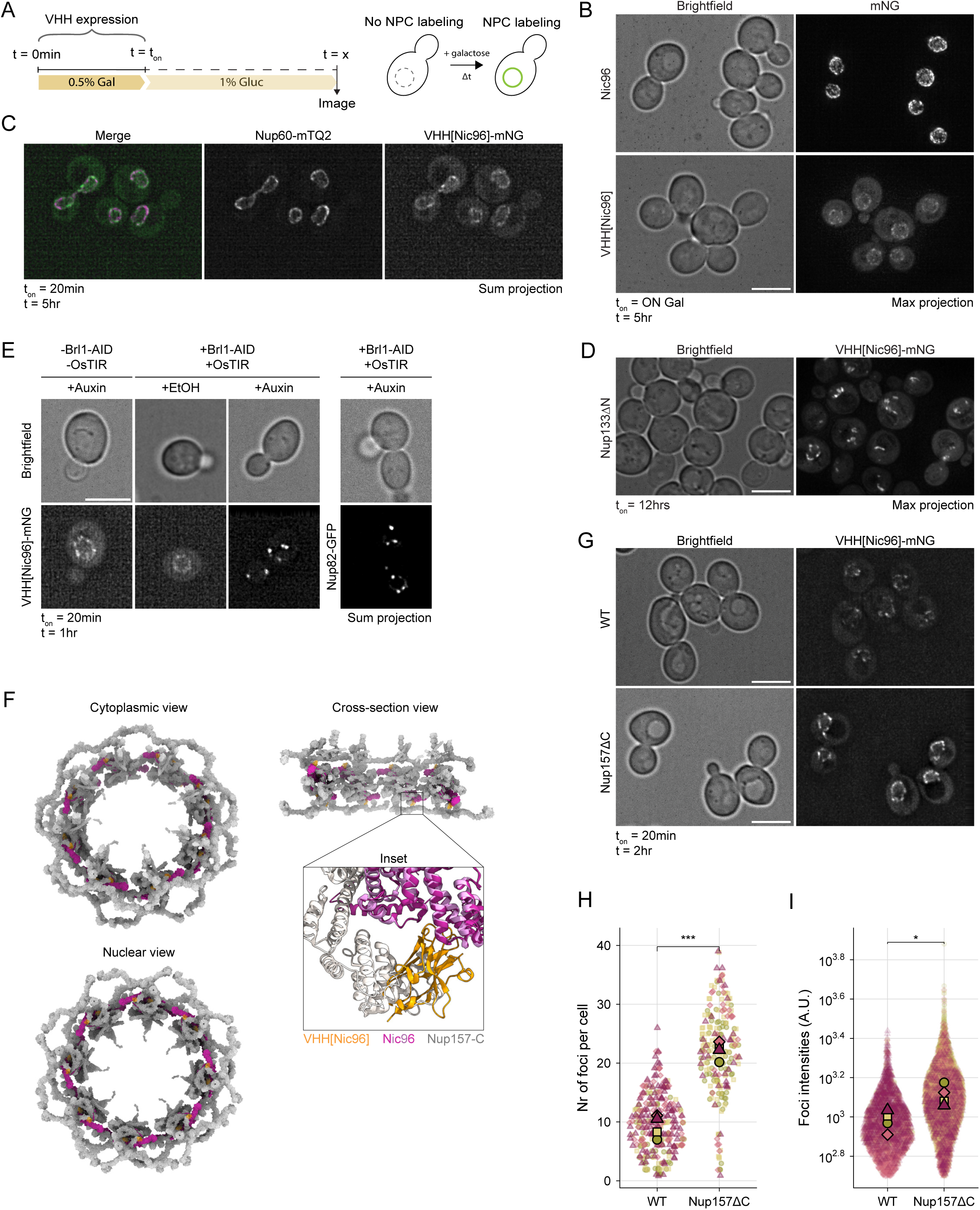
VHH[Nic96]-mNG is incorporated in NPCs during assembly **A**. Experimental setup for imaging experiments. mNeongreen (mNG)-fusions of nanobodies are expressed from an inducible galactose promoter. A short, 20-minute expression burst (t_on_; addition galactose (Gal)) is followed by addition of glucose (Gluc) to stop expression and to allow visualization of nanobody incorporation at the NE over time. Images are taken at various timepoints (t). **B.** Localization of VHH[Nic96]-mNG compared to Nic96-mNG after overnight (ON) induction and a 5-hour glucose chase. Brightness/contrast settings are the same for both panels. Scale bar = 5 µm. Max projection. **C**. Colocalization of VHH[Nic96] with the endogenously tagged Nup Nup60-mTurqoise2 (mTQ2) at t=5hrs following a 20-minute expression burst. Images represent sum projections of 3 z-slices (midplane +/- 0.5 µm). Scale bar = 5 µm. **D.** Localization of VHH[Nic96]-mNG in nup133ΔN cells at t_on_=ON. Images represent max projection. Scale bar = 5 µm. **E.** Localization of VHH[Nic96]-mNG following Brl1 depletion (+Brl1-AID +OsTIR). Strains were grown in medium supplemented with either auxin or ethanol (EtOH) 3hrs prior to VHH[Nic96] induction. VHH[Nic96]-mNG expression was subsequently pulsed for 20 minutes and imaged 1hr after initial induction. Nup82-GFP serves as a control to verify Brl1 depletion. Images represent sum projection of sequential z-slices (midplane +/- 0.5 µm). Scale bar = 5 µm. **F**. Model showing the position of Nic96 (pink) and VHH[Nic96] (orange) in the NPC; based on the NPC structure in ^10^ and the structure of Nic96 bound by VHH[Nic96] ^27^. Inset shows potential clash of VHH (orange) bound to one of the outer copies of Nic96 with the C-terminus of Nup157; based on ^27^ and the inner spoke ring of the yeast NPC ^28^. See also Fig. S2D. **G.** Localization of VHH[Nic96]-mNG in cells with truncated Nup157(aa1-681) (Nup15711C) compared to WT cells at t=2hr following a 20-minute expression burst. Brightness/contrast settings are the same for both panels. Scale bar = 5 µm. Images represent single z-slices. **H.** Quantification of number of VHH[Nic96]-mNG foci per cell in WT cells and Nup15711C cells in panel G using the PunctaFinder plugin (see Fig. 5; see methods). Each dot represents a cell, colours reflect biological replicates. Means of each replica are overlayed. Nr of cells analysed: Nup15711C = 215, WT = 209. Statistical significance was assessed using a negative binomial generalized linear mixed model (GLMM), with predictor effects evaluated using Wald chi-square tests (ANOVA). *** indicates p < 0.001. **I.** As in panel H but now quantifying intensities of VHH[Nic96] signal; each dot represents a focus. A.U. = arbitrary units. Log-transformed normalized intensities were analyzed using a linear mixed-effects model. Statistical significance was assessed using ANOVA with Kenward-Roger degrees of freedom. * indicates p < 0.05, p = 1.86 x 10^-2^.

Next, we assessed if the incorporation of VHH[Nic96] is diminished when we stall NPC assembly. We depleted the essential NPC assembly factor Brl1 using an auxin-inducible degron (AID) system, introducing stalled NPC assembly intermediates in the NE and the mislocalization of late-assembling Nups like Nup82 to cytosolic foci ^23,35^. We depleted Brl1 for 3hrs, then pulsed VHH[Nic96]-mNG expression for a short, 20-minute period and visualized its localization after 40 minutes (Fig. 2E). As previously reported ^23^, Brl1 depletion resulted in the appearance of cytosolic foci of Nup82-GFP under these conditions (Fig. 2E). Importantly, the labelling pattern of VHH[Nic96]-mNG dramatically changed, as we observed a clear clustering when Brl1 is depleted (Fig. 2E). Stalling NPC assembly thus interferes with the incorporation of VHH[Nic96] in NPCs and abolishes the typical NPC-like labelling pattern. We conclude that the incorporation of VHH[Nic96]-mNG in NPCs is dependent on an NPC assembly event.

Lastly, we examined the inaccessibility of the Nic96 epitope in the context of mature NPCs. Crystal structures of VHH[Nic96] bound to the Nic96 C-terminal domain are available ^27^. Superimposing the VHH[Nic96]-bound Nic96 C-terminal domain on the recent high-resolution cryo-EM structure of the inner spoke ring of the yeast NPC ^10,28^ suggests a clash between VHH[Nic96] bound to its epitope and the C-terminal domain of Nup157 and Nup170 (Fig. 2F, view on single spokes in Fig. S2D). To test this experimentally, we made a C-terminal truncation of Nup157, which should resolve the clash between VHH and the inner ring structure, and compared the incorporation of VHH[Nic96] two hours after a short induction pulse with that in wild type cells. Indeed, deleting the C-terminal domain of Nup157 significantly increased the number and intensity of VHH[Nic96]-mNG foci compared to WT cells (Fig. 2GHI). Of note, we could not assess the impact of removing the C-terminal domain from Nup170, as the localization of VHH[Nic96] in the context of both the full deletion and C-terminal deletion of Nup170 is indicative of assembly problems (see Fig. 6HI).

Together, the biochemical and imaging analyses (Fig. 1, 2) show that VHH[Nic96] poorly labels mature NPCs as its epitope is hidden in the context of mature NPCs. Instead, it rapidly and strongly ^27^ binds newly synthesized Nic96, its co-translationally assembled binding partners of the CTN-trimer, Nup192 and Pom152 prior to being incorporated into nascent NPCs.

### VHH[Nic96]-mNG labelling of NPCs does not compromise NPC functionality

We next examined whether the incorporation of VHH[Nic96] impacts NPC biology. Firstly, we assessed if the nuclear transport of a Nab2NLS-mCherry reporter ^36^ was altered upon expression of VHH[Nic96]-mNG. We find that the nuclear accumulation of Nab2NLS-mCherry was not significantly different in wildtype or VHH[Nic96]-mNG expressing strains (Fig. 3A). Secondly, examining the localization pattern of Nup82-mCherry, a Nup that localizes to distinct cytosolic foci in the case of assembly problems ^14,23,37–39^ (Fig. 2E), did not reveal any abnormalities after 6 hours of VHH[Nic96]-mNG overexpression (Fig. 3B).

**Fig. 3.**
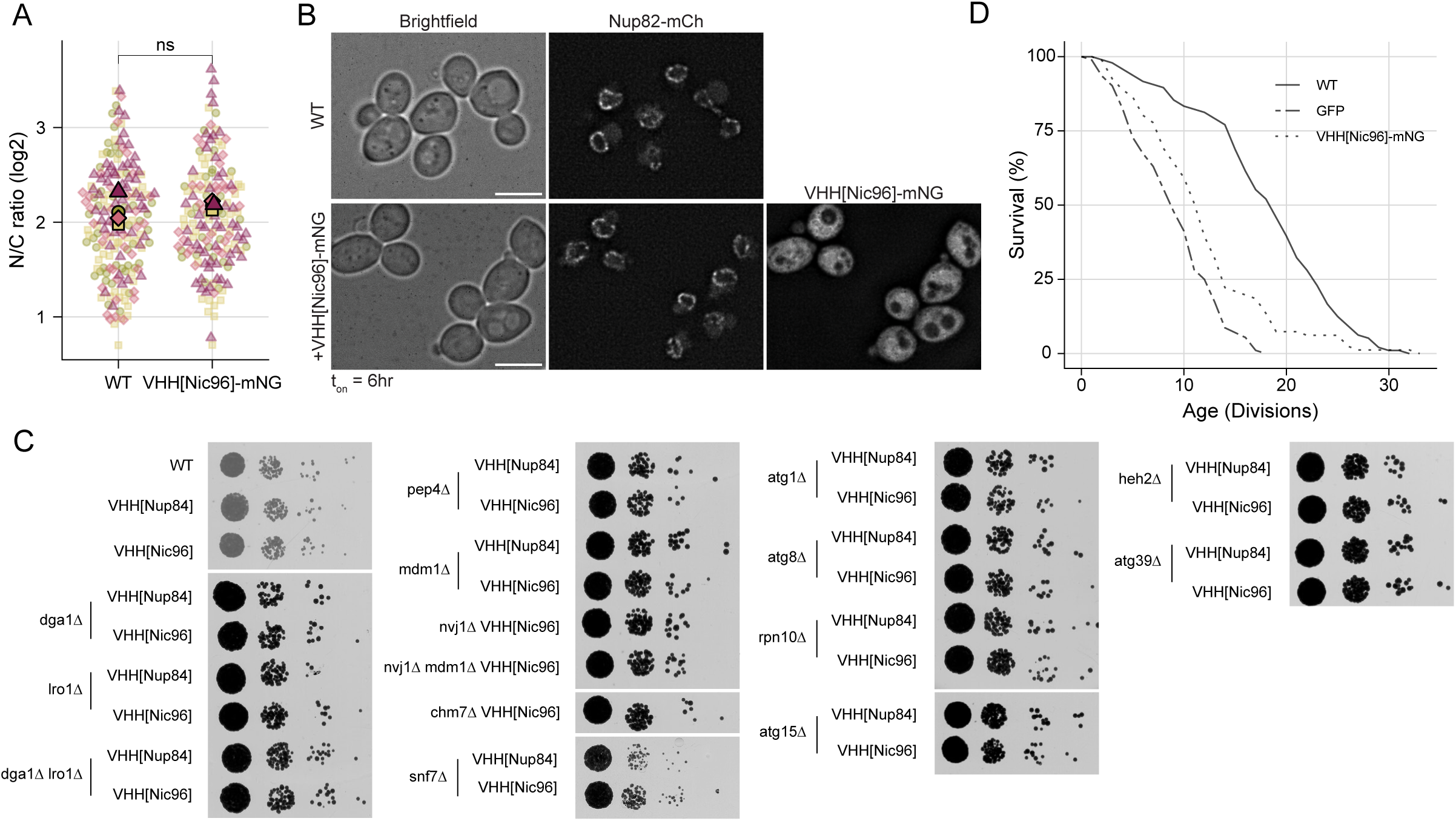
VHH[Nic96]-mNG labelling of NPCs does not compromise NPC functionality **A.** Accumulation of a Nab2*NLS*-mCherry reporter in the nucleus (N) over cytosol (C) - N/C ratio - in wildtype cells and in cells expressing VHH[Nic96]-mNG cells for 3 hours (t_on_). Replica means are overlayed, biological replicas are color-coded. p = 0.5 using a Mann-Whitney test. n > 150 cells from 4 biologically independent experiments. ns = not significant. **B.** Localization of Nup82-mCherry (mCh) in WT cells and cells expressing VHH[Nic96]-mNG after 6 hours on 0.5% galactose (t_on_). Note that in the VHH[Nic96]-mNG channel, high cytosolic backgrounds in absence of a glucose chase period prevent clear delineation of the NE. Images represent single z-slices. Scale bar = 5 µm. **C.** Growth of strains with deletions in indicated NPC quality control pathways expressing VHH[Nup84]-mNG or VHH[Nic96]-mNG compared to WT strain (W303). Columns represent 10-fold serial dilutions, plated on YPGal. **D.** Replicative lifespan analysis of VHH[Nic96]-mNG expressing cells compared to wildtype (WT) cells and wildtype cells expressing GFP, indicating the fraction of cells still alive after x numbers of division.

Thirdly, we reasoned that if VHH[Nic96] would induce assembly problems, abrogating NPC quality control pathways in this background would reveal epistatic interactions. NPC assembly mutants require various quality control pathways to mitigate assembly stress, including lipid droplets, autophagy and the NVJ ^40^. Similarly, the ESCRT pathway has been implicated in NPC quality control ^41^. We made single or double deletions in genes within the autophagy pathway (*atg1*, *atg8*, *atg15* and *atg39*), ESCRT pathway (*chm7*, *heh2, snf7*), lipid droplet metabolism (*lro1*, *dga1*), the NVJ (*nvj1*, *mdm1*) and protein turnover (*pep4*, *rpn10*) to examine their effects on cell viability when combined with the expression of VHH[Nic96]. As a control we combined the same deletions with the expression of VHH[Nup84]. Of note, several deletion strains (*vps4*, *apq12*, the quadruple mutant *lro1 dga1 are1 are2* and *nup116*) easily picked up suppressors during cultivation irrespective of VHH[Nic96] expression and could therefore not be included in our analysis. Importantly, none of these NPC quality control-related genes impacted colony size, indicating that cells do not depend on any of these pathways to grow and divide, even when VHH[Nic96]-mNG is continuously overexpressed (Fig. 3C).

Lastly, we confirmed that growth rates are not affected by overexpression of VHH[Nic96] (Fig. S2E) as previously reported ^27^. However, small differences in cell physiology can become apparent only in the context of ageing. For example, the continuous expression of GFP does not lead to a detectable defect in growth assays but does show a decrease in replicative life span of cells ^42^. To address if the expression of VHH[Nic96] might aggravate the naturally occurring age-related NPC assembly problems ^42^, we determined the replicative life span of cells expressing VHH[Nic96]. The continuous overexpression of VHH[Nic96]-mNG did not shorten lifespan compared to a control strain continuously expressing GFP (Fig. 3D).

Altogether, the absence of defects in transport, growth, and lifespan, as well as the lack of genetic interactions with NPC quality control mutants in VHH[Nic96]-expressing strains, indicates that the incorporation of VHH[Nic96]-mNG into NPCs neither impairs NPC assembly nor compromises NPC function to an extent that results in measurable defects. We envision that the here presented strategy to use nanobody-based labelling of an inaccessible Nup to follow its recruitment to the assembly site can be extended to other inaccessible Nups, although it is worth noting that some epitopes may be more sensitive to misassembly than others, as has been shown *ex vivo* for some xenopus nanobodies ^24^.

### Visualizing VHH[Nic96] shortly after its synthesis provides a view on early assembly

Having thoroughly validated that VHH[Nic96] is incorporated during NPC assembly in living cells, we used it to visualize the early events of incorporation of VHH[Nic96]-mNG-bound subcomplexes into NPCs in more detail. When examining VHH[Nic96]-mNG localization at early time points post a short, 20-minute induction pulse, VHH[Nic96]-mNG localized to the cytosol and in distinct foci at the nuclear periphery (45 min; Fig. 4AB, see Fig. S3A for t=2hr), which, based on our previous characterizations, we expect to represent VHH[Nic96]-bound Nic96 subcomplexes somewhere along the path of incorporation into a new NPC. The homogeneous labelling of NPCs in the NE by VHH[Nup84], which is expressed to similar levels (Fig. S3B), is shown as comparison (Fig. 4AB, see Fig. S3A for t=2hr). Consistent with the lower propensity of VHH[Nic96] to label existing NPCs and the rapid binding of VHH[Nup84] to Nup84 in the context of a fully assembled NPC ^29^, we do not observe a cytosolic VHH[Nup84] pool that is present for VHH[Nic96] (Fig. 4AB). We accidentally noticed a striking similarity of VHH[Nic96] localization to the localization observed for Kap121-GFP (Fig. 4C), which has also been implicated in NPC assembly ^43^, and we indeed find that both overlap (Fig. 4D). The VHH[Nic96]-mNG foci are mobile in the NE and are transmitted to daughter cells during division (Fig. 4E; video S1), indicating they are not mother-retained NPC structures such as SINCs ^41^. We further characterized VHH[Nic96]-mNG foci at higher resolution using stimulated emission depletion (STED) microscopy in living cells and compared them to Nic96-Halo. The signal of VHH[Nic96]-Halo was weaker, but both proteins showed structures with a diameter of ca. 100 nm corresponding to established NPC sizes ^7^, as well as structures of about 200-300 nm which may reflect multiple NPCs in close proximity or the movement of structures along the NE (Fig. 4F, more examples in Fig. S3C). Occasionally, we observed two VHH[Nic96]-Halo structures of approximately 100 nm close to each other (Fig. 4F, Fig. S3C).

**Fig. 4.**
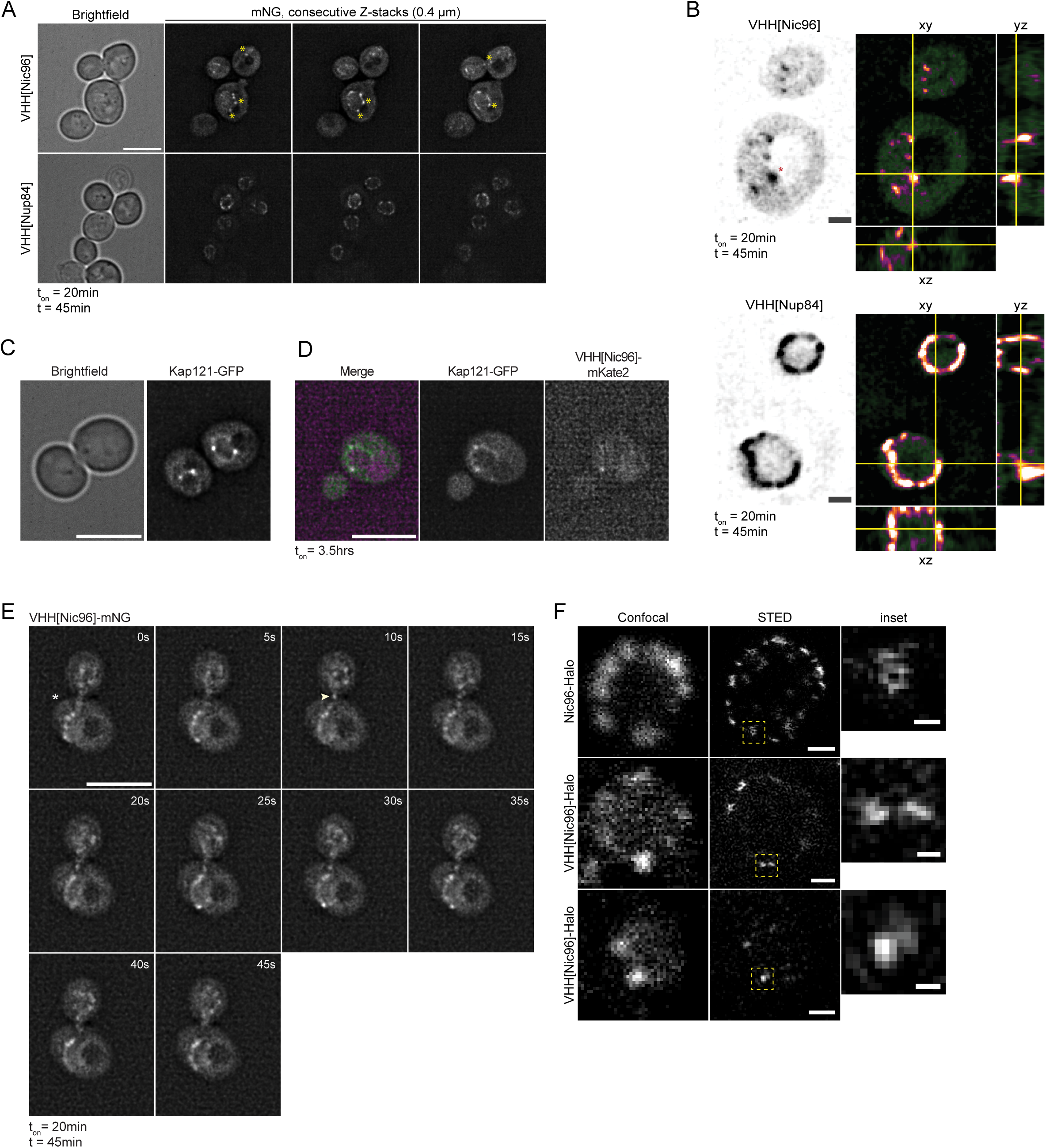
Visualizing VHH[Nic96] shortly after its synthesis **A**. Typical images of the localization of VHH[Nic96]-mNG and VHH[Nup84]-mNG at t=45min following a 20-minute expression pulse. Each panel represents a single z-slice in a consecutive series of z-slices (0.4 um between slices). Brightness/contrast settings are the same for all images and identical to Fig. 5B and Fig. S3A. Asterisk indicate NVJ-proximal focus. Scale bar = 5 µm. **B**. Single z-slice of yeast nuclei depicting distinct foci labelling at the NE by VHH[Nic96]-mNG versus uniform NE labelling by VHH[Nup84]-mNG at t=45min following a 20-minute expression burst. Orthogonal views of the same slice in yz and xz dimensions along yellow lines are depicted with enhanced visibility. Brightness/contrast settings are the same for both images. Asterisk indicate NVJ-proximal focus. Scale bar = 1 µm. **C.** Representative image of cells expressing endogenously tagged Kap121-GFP. Scale bar = 5 µm. **D**. Representative images of Kap121-GFP cells co-expressing VHH[Nic96]-mKate2 for 3.5hrs (t_on_). Panels represent single z-slices. Scale bar = 5 µm. **E.** Time-lapse images in a single z-slice of a dividing cell in which VHH[Nic96]-mNG foci move in the bridge (arrowhead in frame 10s) between mother and daughter cells. Mother cell is indicated by an asterisk. Timestamps indicate time in seconds. Scale bar = 5 µm. See video S1. **F**. Live-cell confocal and STED microscopy images of Nic96-Halo and VHH[Nic96]-Halo structures in the yeast nuclear envelope labelled with SiR-Halo dye. VHH[Nic96]-Halo expressing cells were imaged at t=1hr following a 20-minute expression burst. All STED images are processed using a Wiener filter. Scale bar = 0.5 µm for the confocal and STED images and scale bar = 0.1 µm for inset.

Next, we asked if a quantitative assessment of the VHH[Nic96]-mNG signal might be used to assess rates of assembly. We used the PunctaFinder plugin ^44^, that, based on calculated threshold values (Fig. 5A, see methods), reports foci numbers and their intensity in 3D in individual yeast cells (Fig. 5B, see Fig. S4A for intensities of detected foci). With the established threshold, which we set to minimize false positive candidates, PunctaFinder underestimates the true number of VHH[Nic96] foci per cell. Furthermore, given the inherent biological variation in the short expression burst from the gal expression system and the rate of cell division, the absolute number and intensity of VHH[Nic96]-mNG foci are somewhat variable between experiments. In spite of this, we consistently found that following the 20-minute expression burst, the number and intensity of VHH[Nic96]-mNG foci increased in the first hours and then decreased again as cellular VHH[Nic96] levels decrease and labelled NPCs dilute out to daughter cells (Fig. 5CD, see S4D for total cell intensities). The number of foci approaches that of constitutively expressed VHH[Nic96]-mNG (Fig. S4E). Based on a comparison to endogenously tagged Nic96-mNG, we estimate that on average half of the Nic96 copies within a focus are bound by VHH[Nic96] (Fig. 5DE). We next quantified the VHH[Nic96]-mNG foci in haploid and diploid cells after a short expression pulse and fitted the data with a generalized linear mixed model (see methods). Although more frequent sampling would be required for a precise quantification of the time difference, the model predicts that in diploid cells the maximum number of VHH[Nic96] foci is reached earlier than in haploid cells (Δt_max_ = 1.39 hr; 95% CI [1.26, 1.52]), which suggests a higher rate of NPC assembly (Fig. 5F). This is consistent with our estimates that diploid cells have approximately twice as many NPCs (based on cell- and nuclear size ^45^ and relative abundances of Nups ^46^) while dividing with similar division times (Fig. S5A; ^47^) Together, the quantitative characterization of the VHH[Nic96]-mNG signal is consistent with the continuous synthesis of NPCs throughout the cell cycle ^13^ and the dilution of NPCs to daughter cells and suggests increased assembly rates in diploid cells compared to haploid cells.

**Fig. 5.**
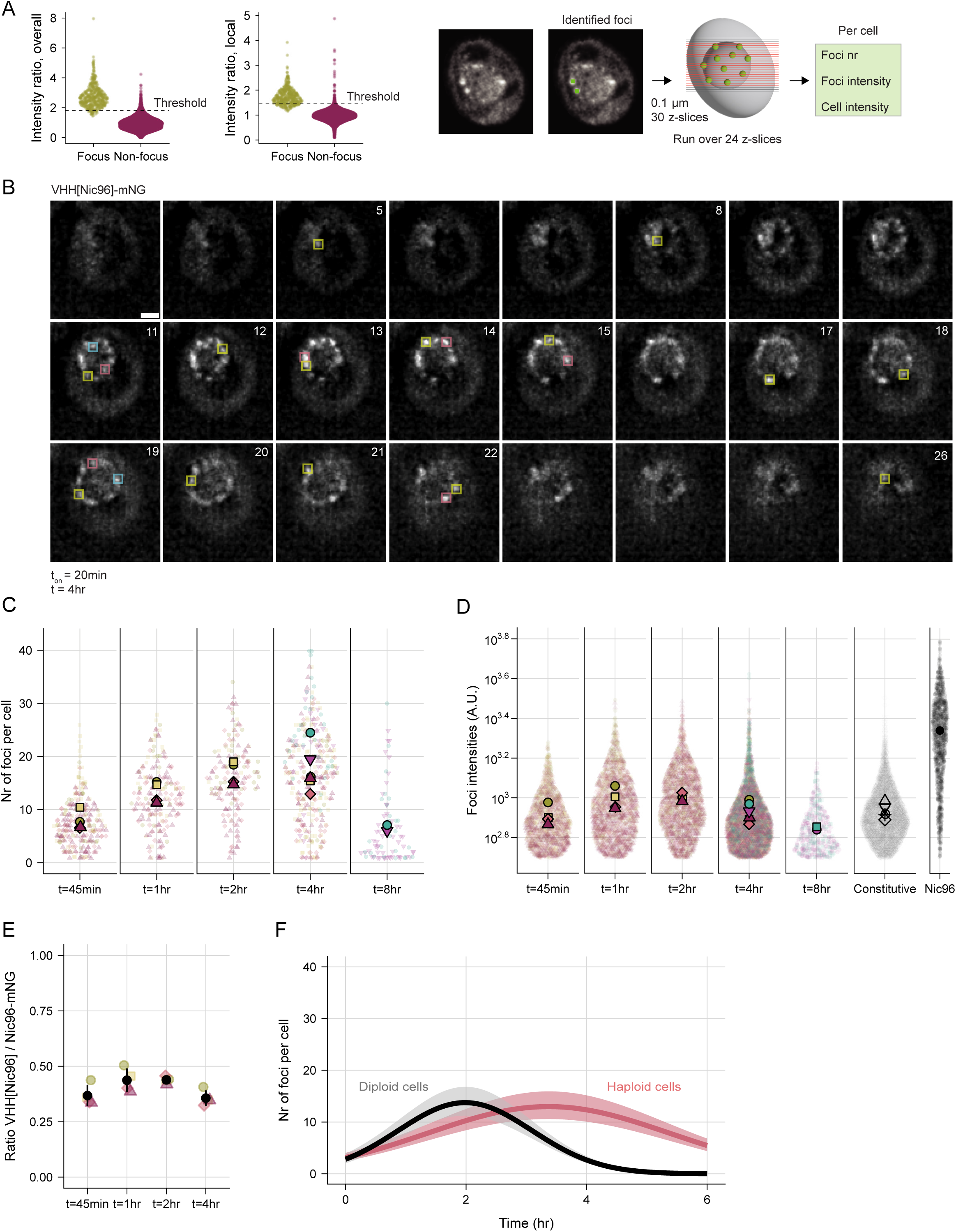
Quantitative assessment of VHH[Nic96] labelling is consistent with dynamics of assembly and cell division **A.** Intensity ratios determined by PunctaFinder for punctate (focus) and nonpunctate regions (non-focus). Thresholds values (dashed lines) were determined by bootstrapping (see methods); n = 453 visually detected foci in 91 images. Z-stacks of 24 images with a separation of 0.1 µm (depicted by red lines, the first 3 and last 3 z-stacks are excluded from analysis) were analysed with PunctaFinder to report the total number of foci per cell, their intensity and the average cell intensity. An example of identified foci in a single slice of a z-stack is given, where the two identified foci (5×5 pixel area) are depicted in green; the other visible foci are identified in different z-slices where they reach their maximum. **B.** Example of VHH[Nic96]-mNG foci detected in a single cell at t=4hr following a 20-minute expression burst. Coloured squares indicate detected foci (5×5 pixels). See Fig. S4A for foci intensities. Images represent single z-slices. Numbers indicate Z-slice. Scale bar = 1 µm. **C.** Number of foci per cell at each sampling timepoint. Each dot represents a cell, colours reflect independent experiments. Replica means are overlayed. Each timepoint up to 4 hours contains data from at least four independent experiments (see Fig. S4B for individual replicas) and >140 cells. The 8hr timepoint contains data from 2 independent experiments (n=81 cells) (see Fig. S4C for individual replicas). **D.** Foci intensities when comparing VHH[Nic96] expressed from galactose promoter (same dataset as in C; see Fig. S4F for individual replicas) to VHH[Nic96] expressed from a Nup120 promoter (constitutive, >40 cells per replica) and to endogenous Nic96-mNG (40 cells). Replica means are overlayed. Each dot represents a single focus. A.U. = arbitrary units. **E.** Ratio between median VHH[Nic96]-mNG focus intensity and median Nic96-mNG focus intensity at each timepoint. Colours indicate the biological replicas; black dots represent mean +/- standard error. **F.** Nr of foci per cell in time following the 20-minute expression burst under the statistical model described in Supplementary Table 1 and Fig. S5DEF for haploid (pink) and diploid (grey) cells. The thick line represents mean values; the coloured band indicates 95% confidence interval. See Fig. S5 BC for individual replicas.

### Association of Nic96 subcomplexes with NE-localized lipid droplets increases when assembly is delayed

Excitingly, our findings suggest that we can visualize and quantify early events of incorporation of VHH[Nic96]-mNG-bound subcomplexes into NPCs, paving the way for detailed spatiotemporal studies of NPC assembly. We set out to further examine the VHH[Nic96] foci that appear close to the vacuole (asterisk in Fig. 4AB, see Fig. S2A for constitutive expression). To better understand the nature of these vacuole-adjacent VHH[Nic96] foci, we resorted to correlative light and electron microscopy (CLEM) and tomography, using Vph1 as a vacuolar marker. Consistent with the incorporation of VHH[Nic96]-mNG in NPCs, the fluorescent signal was detected at an NPC (Fig. 6A top panel, NPC indicated by asterisk). Interestingly, we also found VHH[Nic96]-mNG signal overlapping with lipid droplets (Fig. 6A middle and bottom panel) that were often close to an NPC (Fig. 6B). The general lipid droplet marker Erg6-GFP, as well as Pdr16-GFP and Ldo16-GFP which define a metabolically regulated lipid droplet subpopulation associated with the nucleus-vacuole junction (NVJ) ^48^, colocalized with VHH[Nic96]-mKate2 foci in more than 50% of cases (Fig. 6CD; please note compared to mNG, mKate2-tagged VHH[Nic96] shows only few foci per cell, which likely relates to low fluorescence and longer maturation times ^32^). These findings collectively suggest that newly synthesized Nic96 can be incorporated into nascent NPCs, as well as deposited on lipid droplets in the NVJ area. The association with lipid droplets does not appear to be an essential step towards NPC assembly as VHH[Nic96]-mNG foci at the NE also appear in a strain that lacks lipid droplets (*are1Δ, are2Δ, dga1Δ, lro111* ^49^) (Fig. 6E). Of note, the proximity to lipid droplets visualized by AUTODOT^TM^ is not restricted to VHH[Nic96]-mNG foci within the NVJ area, nor limited to the highest-intensity foci population (arrowheads in Fig. 6F).

**Fig. 6.**
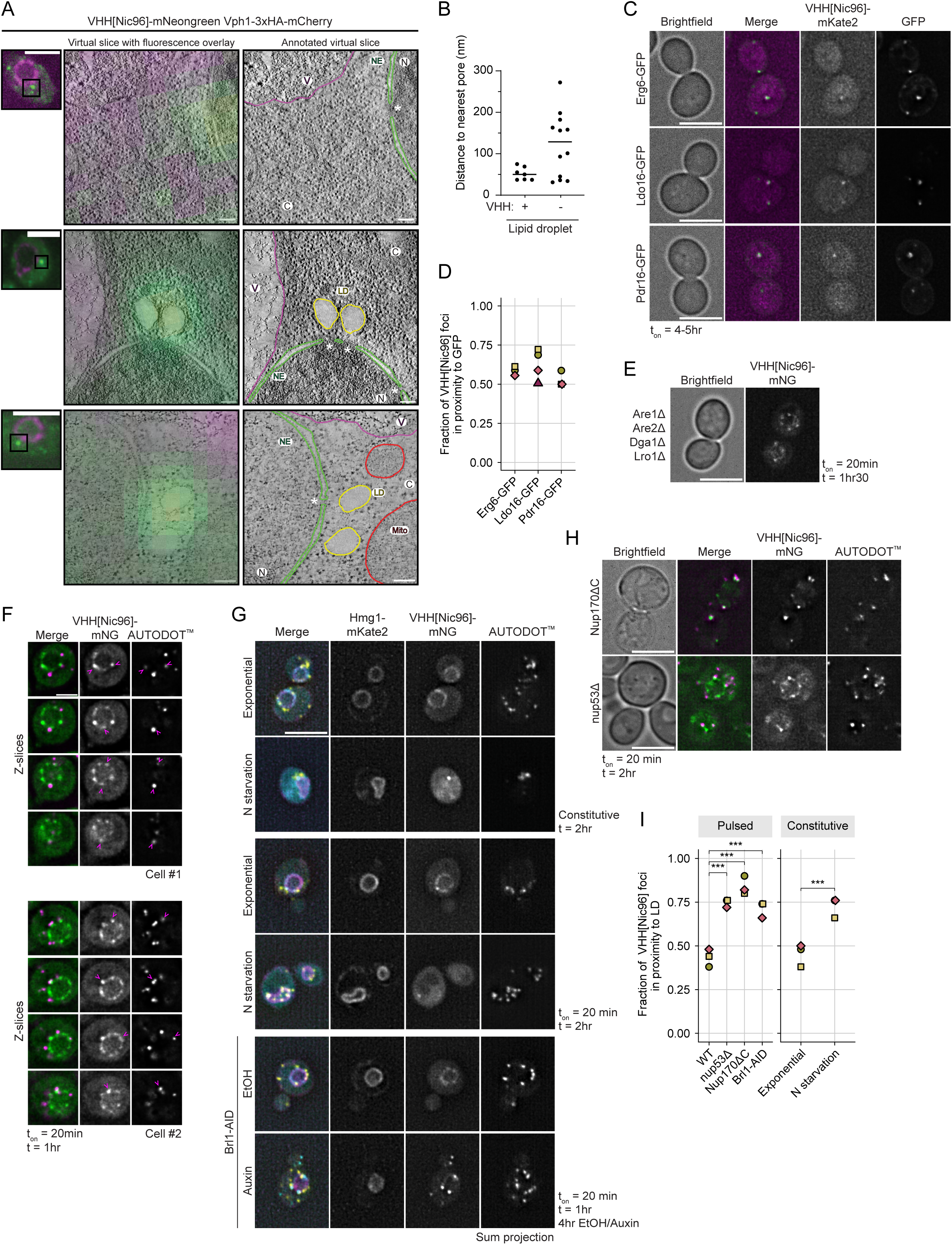
Association of Nic96 subcomplexes with NE-localized lipid droplets **A.** CLEM image showing the overlap of NVJ-proximal VHH[Nic96]-mNG foci with a NPC (top panel) and lipid droplets (bottom 2 panels). Vph1-3xHA-mCherry is used as a vacuolar marker. Insets show fluorescent image; middle panel shows overlay of fluorescence with EM image. Right panel represents annotated slices. LD (yellow) = lipid droplet, NE (green)= nuclear envelope, C = cytosol, N = nucleus, V = vacuole (pink), Mito (red) = mitochondrion. Asterisks indicate NPCs. Scale bar = 100 nm for EM image and 3 µm for inset. **B.** Quantification of distance to nearest NPC for LDs with and without VHH[Nic96]-mNG signal. Line indicates mean distance. **C.** Examples of VHH[Nic96]-mKate2 foci overlapping with Erg6-GFP, Ldo16-GFP and Pdr16-GFP. VHH[Nic96]-mKate2 expression was induced for 4-5hrs (t_on_). Each panel represents a single z-slice. Scale bar = 5 µm. **D.** Quantification of the fraction of VHH[Nic96]-mKate2 foci in proximity to GFP signal. Each point represents a biological replicate. Nr of VHH[Nic96]-mKate2 foci analysed in each replica: *34, 62, 72* (Erg6-GFP)*; 83, 36, 51, 87* (Ldo16-GFP); *46, 56, 42* (Pdr16-GFP). **E.** Localization of VHH[Nic96]-mNG in a lipid droplet dead strain (*are111, are211, dga111, lro111*). Images were taken at 1h30 following a 20-minute expression burst and represent a single Z-slice. Scale bar = 5 µm. **F.** Typical images of individual cells showing NE-localized VHH[Nic96]-mNG foci in multiple z-slices close to lipid droplets co-stained with the lipid droplet dye AUTODOT^TM^ at t=1hr following a 20-minute expression burst. Each panel represents a single z-slice. Arrowheads point to VHH[Nic96]-mNG foci near or overlapping with lipid droplets. Scale bar = 2 µm. **G.** Typical images of VHH[Nic96] localization after nitrogen starvation and Brl1-depletion co-stained with the lipid droplet dye AUTODOT^TM^. Hmg1-mKate2 is used as a NE marker. Cells were starved for nitrogen for 2hrs and VHH[Nic96]-mNG was either constitutively expressed from a Nup120 promoter or pulsed for 20 minutes concomitant with N starvation. Brl1 was depleted as in Fig. 2E, VHH[Nic96] expression was pulsed for 20 minutes in the last hour. Images represent sum projection of sequential z-slices (midplane +/- 0.5 µm). Scale bar = 5 µm. **H.** Examples of VHH[Nic96]-mNG localization in NPC mutant strains co-stained with the lipid droplet dye AUTODOT^TM^ at t=2hr following a 20-minute expression burst. Each panel represents a single z-slice. Scale bar = 5 µm. **I.** Quantification of the fraction of VHH[Nic96]-mNG foci in proximity to lipid droplets in various conditions shown in panels F, G and H. For each replicate, 50 highest intensity foci were scored for LD proximity. Each point represents a biological replicate. Separate binomial generalized linear models were fitted to the pulsed and constitutive datasets, followed by post hoc analysis using estimated marginal means (emmeans) with planned pairwise comparisons to respective control (trt.vs.ctrl; ctrl = WT (pulsed) or exponential (constitutive)). *** indicates p < 0.0001.

The deposition of VHH[Nic96] on lipid droplets close to the NE may reflect the storage of Nups that cannot be accommodated at the NE, as suggested in an earlier study ^50^, and our data implies that the pathway to deposit Nups on lipid droplets involves, at minimum, newly synthesized NPC building blocks. To further examine the relation between Nic96 subcomplex assembly and Nic96 subcomplex association with lipid droplets, we assessed VHH[Nic96] localization in contexts that may shift the balance from assembly towards storage, namely starvation and impaired assembly. In exponentially growing WT cells, approximately 40-50% of the brightest VHH[Nic96]-mNG foci at the NE were in close proximity to lipid droplets visualized by AUTODOT^TM^ (arrowheads in Fig. 6F), both after pulsed expression and when constitutively expressed (Fig. 6I). Upon nitrogen starvation, this frequency increased to about 75% (Fig. 6I), and, consistent with a delay in VHH[Nic96] incorporation in NPCs, VHH[Nic96] notably changed its localization (Fig. 6G, upper and middle panel). In nitrogen-starved cells, the NE showed only few foci instead of the typical NPC-like labelling pattern, and the cytosolic VHH pool increased (quantified in Fig. S6A). When we expressed VHH[Nic96]-mNG in backgrounds where we depleted Brl1 ^23^ (Fig. 6G, bottom panels) or where the NPC structure and/or assembly is compromised, such as deletions of Nup53 and the C-terminus of Nup170 (Fig. 6H), VHH[Nic96]-mNG foci were found more frequently in proximity to lipid droplets (Fig. 6I). Notably, in all three cases, a population of higher intensity VHH[Nic96]-mNG foci appears (Fig. S6B), indeed indicating local accumulation of VHH[Nic96]-bound subcomplexes consistent with impaired assembly. Moreover, when we examined lipid droplet proximity of VHH[Nic96] foci in relation to their subcellular localization in Brl1-depleted cells, NE-localized VHH[Nic96] foci tended to be more frequently in proximity to a lipid droplet than cytosolic VHH[Nic96] foci (Fig. S6C). Thus, in all mutants and conditions, and irrespective of whether VHH[Nic96] was pulsed or constitutively expressed, we observed an increase in the fraction of lipid droplet-proximal VHH[Nic96] foci when interfering with assembly (Fig. 6I), indeed indicating a coupling between efficiency of NPC biogenesis and association of Nic96 subcomplexes with NE-localized lipid droplets.

Collectively, these findings support a model (Fig. S6D) in which new Nic96 subcomplexes are incorporated into new NPCs or stored on lipid droplets, that are predominantly at the NE and close to the assembly site. Our data suggests that in conditions of cellular stress, the spatial association between lipid droplets and NPC assembly intermediates increases, which may provide a mechanism to temporarily stall unassembled or misassembled subcomplexes on lipid droplets close to the assembly site and/or promote their degradation.

## Discussion

The assembly and quality control of nuclear pore complexes (NPCs) are crucial processes for all eukaryotic life. Addressing the many unresolved questions regarding the assembly process — such as the biochemical landscape of NE regions that permit NPC biogenesis, and the factors involved in timing and guarding the assembly process — requires tools for visualization and biochemical tracking of assembling NPCs. The here reported probe opens avenues of spatiotemporal characterization of NPC assembly and the associated quality control pathways.

In this work, we show that a nanobody against Nic96, VHH[Nic96] ^27^, can be used to report on NPC assembly events in living yeast cells. The inner ring component Nic96 is buried deep within the structure of mature NPCs ^28^ (Fig. 2F), thereby decreasing the likelihood of VHH[Nic96] binding within existing structures and biasing VHH[Nic96] labelling towards accessible Nic96 copies in the pre-assembly and assembly stages (Fig. 1). Indeed, our biochemical and live-cell imaging data support that VHH[Nic96] becomes part of the Nic96-Nsp1-Nup49-Nup57 subcomplex and the Nic96-Nup192 inner ring connection and depends on an assembly event to be incorporated in NPCs (Fig. 1, 2). Incorporation of VHH[Nic96] can be used to assess the rate of assembly (Fig. 5). Fortunately, VHH[Nic96]-mNG labelling of NPCs does not compromise NPC functionality (Fig. 3). Illustrating that the nanobody is a unique tool to address NPCs assembly with temporal and spatial resolution in living cells, our findings highlight three insights that present exciting avenues for future research. First, we report on newly synthesized Nic96 subcomplexes residing close to lipid droplets at the NE, and more so in conditions when assembly is delayed (Fig. 6). Our CLEM data shows examples of VHH[Nic96]-bound subcomplexes deposited on lipid droplets in the NVJ area, which implies storage pools of unassembled subcomplexes close to the assembly site. Second, the overlap of Kap121-GFP foci with VHH[Nic96] foci (Fig. 4BC) strongly suggests their co-recruitment to the same NE region and provides a starting point for further determining the biochemical landscape of the assembly site. VHH[Nic96] thus provides a valuable tool to assess assembly dynamics and complements biochemical, *in vivo* and *ex vivo* strategies that have been crucial in determining the sequential order of NPC assembly ^16,22,24–26,51^

The finding that in exponentially growing wild type cells, newly synthesized Nic96 subcomplexes associate with lipid droplets (Fig. 6) raises interesting questions. As VHH[Nic96] is still incorporated in NPCs in mutant strains lacking lipid droplets, we conclude that lipid droplets are not an intermediate in NPC assembly but rather function as storage pools. We reason that under non-stress conditions, transient sequestration of Nups on lipid droplets may provide a buffered supply of NPC building blocks. The association of nanobody-labelled new Nups with Pdr16- and Ldo16-positive lipid droplets near the NVJ could reflect a stable and local Nup storage platform, as exponentially growing cells always display at least one Pdr16-positive lipid droplet ^48,52^ and this lipid droplet subpopulation has been shown to be resistant to lipolysis ^52,53^. Whether this lipid droplet subpopulation actively contributes to Nup regulation is an interesting question that follows from this work. Storage pools of unassembled subcomplexes may be especially critical when NPC assembly failure is more frequent, as in NPC assembly mutants. In support of this, lipid droplets and the NVJ are required to alleviate NPC assembly stress through yet unclear mechanisms ^40^. Additionally, strains incapable of forming lipid droplets show an extended lag phase ^54^, which might in part be attributed to the lack of such Nup storage pools, necessitating the *de novo* synthesis of Nups for NPC assembly upon re-entry into the cell cycle.

Multiple Nups, amongst others Nic96, Nup57, Nup82, Nup192, Pom152 and the nuclear transport receptor Kap121, have been found on lipid droplets under conditions of metabolic or ER stress ^50,55^. For several Nups, lipid-binding plays a role in NPC biogenesis ^56^ and the idea that lipid droplet dynamics contribute to NPC homeostasis by buffering excess Nups in stress conditions has been proposed before ^50^. Our findings extend this model by demonstrating that newly synthesized Nups are deposited on these droplets. Additionally, as we can visualize their subcellular localization, our findings suggest this deposition occurs in proximity to the assembly site, as we show that the lipid droplets engaging in these interactions are predominantly at the NE. We speculate that depositing new Nic96 subcomplexes on lipid droplets close to the assembly site might be a way to maintain low levels of cytosolic, unassembled Nups when the incorporation into new NPCs is compromised. Maintaining low levels of specifically FG-Nups is likely important to prevent their condensation and toxic aggregation, as the FG-Nups readily form condensates and aggregates outside of the context of the NPC ^18,19,57,58^. Furthermore, cytosolic Nup levels are implicated as a regulator of assembly initiation ^59,60^. Their deposition on lipid droplets thus conceivably helps in preventing novel assembly events in conditions of stress and may even facilitate their degradation, as has previously been shown for Nup159 under conditions of ER stress ^55^. How the interaction between lipid droplets and new Nic96 subcomplexes is orchestrated, will be an exciting future avenue. The interactions with lipid droplets might be regulated through Nic96 itself as it contains a predicted fatty acid (FAA) binding domain that proposedly drives the release of Nic96 from lipid droplets during lipolysis ^50^.

We were intrigued that even in unperturbed growth conditions we find such frequent proximity of new Nic96 subcomplexes to lipid droplets (Fig. 6I). We consider two possible scenarios. First, lipid droplet proximity might reflect local storage or turnover of un- or misassembled Nic96 subcomplexes. In this view, lipid droplets act as a buffering system to sequester aberrant or excess subcomplexes and/or lipids that arise during NPC (mis)assembly. In addition, misassembly rates in exponentially growing cells are currently unknown, thus raising the possibility that VHH[Nic96] visualizes baseline rates of un- or misassembled subcomplex turnover. Several observations are consistent with this hypothesis: backgrounds that display elevated NPC misassembly events, including ageing ^42^ and NPC assembly mutants ^40,61^, display increased numbers of lipid droplets, and in similar misassembly contexts we find VHH[Nic96] more often adjacent to a lipid droplet (Fig. 6I). Alternatively, one may speculate that lipid droplets are spatially linked to NPC biogenesis, where NE regions that support lipid droplet formation may also be intrinsically permissive for NPC assembly, or vice versa. Computational models of the oligomeric structure of yeast Brl1/Brr6 ^35^ and metazoan CLCC1 ^62^, proteins that contribute to membrane fusion during NPC biogenesis ^35,62,63^, predict a channel suited for lipid remodelling at the inner and outer nuclear membrane fusion site ^35,62^. Whether such local lipid remodelling processes contribute to lipid droplet formation, or whether - vice versa - local membrane remodelling and lipid flow preceding lipid droplet formation could contribute to establishing the fusion site of the two NE leaflets is an interesting future endeavour. Similarly, close proximity of lipid droplet-resident proteins such as lipid metabolism enzymes and/or lipid droplet-mediated lipid exchange may conceivably contribute to membrane remodelling required for NPC biogenesis.

The proximity of new NPCs to lipid droplets specifically in the NVJ-area (Fig. 6A) under unperturbed growth conditions makes it tempting to speculate that NPC synthesis may preferentially occur in this NE region, raising the possibility that the role previously attributed to this region in handling misassembled NPC intermediates ^40^ reflects a more general pathway that becomes particularly apparent under conditions that challenge NPC assembly. Lipid droplet organization in the NVJ region, including the coordination of lipid droplet–vacuole contact sites, is tightly coupled to cellular metabolic state ^64,65^, and these lipid droplets have been proposed to be well positioned to mediate localized protein removal ^65^. It is thus tempting to speculate that this environment could support NPC biogenesis by coupling assembly with local quality control, thereby facilitating both the incorporation of newly synthesized components and the sequestration or degradation of unassembled subcomplexes.

Collectively, our findings illustrate how the strategy to pulse-label NPC assembly intermediates opens new avenues for dissecting the spatiotemporal regulation of NPC assembly.

## Methods

### Strains and yeast growth conditions

All yeast strains and plasmids used in this study are listed in table 1 and 2. Plasmids were generated by standard molecular biology techniques and validated by sequencing. Yeast strains were validated by microscopy and/or PCR. Please note that we removed a nuclear localization signal that was present in some of the original constructs ^27^; specifically, when generating pAV44 from the TUS4991 plasmid, the SV40-NLS directly after the mKate2 was removed.

**Table 1:** Yeast strains.

| Strain name | Genotype | Description | Source |
| --- | --- | --- | --- |
| W303 | *MATa leu2-3,112 trp1-1 can1-100 ura3-1 ade2-1 his3-11,15* | Wildtype strain | Open biosystems |
| BY4741 | *MATa his3Δ1 leu2Δ0 met15Δ0 ura3Δ0* | Wildtype strain | Open biosystems |
| yAV108 | *W303; leu2::pGal1-VHH[Nup84]-mNeogreen::LEU2* | Strain expressing VHH[Nup84]-mNeogreen from an inducible galactose promoter, integrated in leu2 locus using pAV49 | This study |
| yAV110 | *W303; leu2::pGal1-VHH[Nic96]-mNeogreen::LEU2* | Strain expressing VHH[Nic96]-mNeogreen from an inducible galactose promoter, integrated in leu2 locus using pAV50 | This study |
| yAV235 | *W303; leu2::pNup120-VHH[Nic96]-mNeogreen::LEU2* | Strain expressing VHH[Nic96]-mNeogreen from a Nup120 promoter, integrated in leu2 locus using pAV70 | This study |
| yAV232 | *W303; nup133(301-1157)-PrA::HIS leu2::pGal1-VHH[Nic96]-mNeogreen::LEU2* | Genomic integration of VHH[Nic96]-mNeogreen under inducible galactose promoter using pAV50 in cells expressing a N-terminal truncation of Nup133 | This study |
| yAV189 | *W303; leu2::pGal1-VHH[Nic96]-HALO::LEU2* | Strain expressing VHH[Nic96]-HALO from an inducible galactose promoter, integrated in leu2 locus using pAV64 | This study |
| yAV224 | *W303; Nic96-HALO::Hyg* | C-terminal tagging of Nic96-HALO using pAV61 | This study |
| yAV255 | *W303; Pom34-S-TEV-ZZ::HIS3; leu2::pNup120-VHH[Nic96]-mNeogreen-1Myc7His::KanMX, LEU2* | Pom34-S-TEV-ZZ tagged in its endogenous locus using pKW803; constitutively expressed pNup120-VHH[Nic96]-mNeogreen-1Myc7His integrated in leu2 locus using pAV50 and pYM46 | This study |
| KWY12172 | *W303; his3::pNup120-VHH[Nic96]-S-TEV-ZZ::HIS; lys2Δ::KanMX* | Constitutively expressed pNup120-VHH[Nic96]-S-TEV-ZZ integrated in his3 locus | This study |
| yAV213 | *yAV108; lys2Δ::KanMX, leu2::pGal1-VHH[Nup84]-S-TEV-ZZ::HIS3, LEU2* | VHH[Nup84]-S-TEV-ZZ under galactose promoter integrated in the leu2 locus with His3 selection marker | This study |
| yAV214 | *yAV110; lys2Δ::KanMX, leu2::pGal1-VHH[Nic96]-S-TEV-ZZ::HIS3, LEU2* | VHH[Nic96]-S-TEV-ZZ under galactose promoter integrated in the Leu2 locus with HIS3 selection marker, using pKW803 to replace mNG with S-TEV-ZZ.<br>Lys2 deletion by PCR amplification of lys2 deletion cassette from YKO collection | This study |
| yAV229 | *BY4741; his3::pGal1-VHH[Nic96]-mNeogreen::HIS3* | BY strain expressing VHH[Nic96]-mNeogreen from an inducible galactose promoter, integrated in His locus using pAV69 | This study |
| yAV236 | *BY4741; Nab2NLS-mCherry::LEU* | Expression of a Nab2NLS-mCherry reporter from a centromeric plasmid (pBT014) | This study |
| yAV237 | yAV229; *Nab2NLS-mCherry::LEU* | Expression of a Nab2NLS-mCherry reporter from a centromeric plasmid (pBT014) | This study |
| yAV193 | W303; *nup82-mCherry::NAT leu2::pGal1-VHH[Nic96]-mNeogreen::LEU2* | Genomic integration of VHH[Nic96]-mNeogreen under inducible galactose promoter in Leu2 locus using pAV50 in cells expressing Nup82-mCherry (KWY6765; gift from Weis lab) | This study |
| yAV257 | yAV110; *chm7Δ::Hyg* | Deletion of *chm7* using pFA6-hphNT1 | This study |
| yAV201 | yAV110; *dga1Δ::Hyg* | Deletion of *dga1* using pFA6-hphNT1 | This study |
| yAV200 | yAV110; *lro1Δ::Hyg* | Deletion of *lro1* using pFA6-hphNT1 | This study |
| yAV220 | yAV200; *dga1Δ::Nat* | Deletion of *dga1* using pFA6-NatNT2 | This study |
| yAV167 | yAV121; *heh2Δ::Hyg* | Deletion of *heh2* using pFA6-hphNT1 | This study |
| yAV165 | yAV121; *atg39Δ::Hyg* | Deletion of *atg39* using pFA6-hphNT1 | This study |
| yAV203 | yAV110; *atg1Δ::Hyg* | Deletion of *atg1* using pFA6-hphNT1 | This study |
| yAV130 | W303; *atg8Δ::KanMX leu2::pGal1-VHH[Nic96]-mNeogreen::LEU2* | DTCPL954 strain (gift from Lusk lab) expressing VHH[Nic96]-mNeogreen from an inducible galactose promoter, integrated in LEU locus using pAV50 | This study |
| yAV204 | yAV110; *rpn10Δ::Hyg* | Deletion of *rpn10* using pFA6-hphNT1 | This study |
| yAV158 | yAV121; *pep4Δ::Hyg* | Deletion of *pep4* using pFA6-hphNT1 | This study |
| yAV160 | yAV121; *mdm1Δ::Hyg* | Deletion of *mdm1* using pFA6-hphNT1 | This study |
| yAV159 | yAV121; *nvj1Δ::Hyg* | Deletion of *nvj1* using pFA6-hphNT1 | This study |
| yAV181 | yAV159; *mdm1Δ::Nat* | Deletion of *mdm1* using pFA6-NatNT2 | This study |
| yAV152 | yAV117; *dga1Δ::Hyg* | Deletion of *dga1* using pFA6-hphNT1 | This study |
| yAV151 | yAV117; *lro1Δ::Hyg* | Deletion of *lro1* using pFA6-hphNT1 | This study |
| yAV182 | yAV151; *dga1Δ::Nat* | Deletion of *dga1* using pFA6-natNT2 | This study |
| yAV157 | yAV117; *heh2Δ::Hyg* | Deletion of *heh2* using pFA6-hphNT1 | This study |
| yAV155 | yAV117; *atg39Δ::Hyg* | Deletion of *atg39* using pFA6-hphNT1 | This study |
| yAV153 | yAV117; *atg1Δ::Hyg* | Deletion of *atg1* using pFA6-hphNT1 | This study |
| yAV144 | W303; *atg8Δ::KanMX leu2::pGal1-VHH[Nup84]-mNeogreen::LEU2* | DTCPL954 strain (gift from Lusk lab) expressing VHH[Nup84]-mNeogreen from an inducible galactose promoter, integrated in LEU locus using pAV49 | This study |
| yAV156 | yAV117; *rpn10Δ::Hyg* | Deletion of *rpn10* using pFA6-hphNT1 | This study |
| yAV148 | yAV117; *pep4Δ::Hyg* | Deletion of *pep4* using pFA6-hphNT1 | This study |
| yAV150 | yAV117; *mdm1Δ::Hyg* | Deletion of *mdm1* using pFA6-hphNT1 | This study |
| yAV142 | W303; *nic96-mNeogreen::URA3* | C-terminal tagging of Nic96-mNeogreen using pAV54 | This study |
| yAV186 | yAV110; *nup60-mTQ2::URA3* | C-terminal tagging of Nup60-mTQ2 using pSJ1458 | This study |
| yAV177 | yAV110; *nup170Δ::Hyg* | Deletion of Nup170 using pFA6-hphNT1 | This study |
| yAV179 | yAV110; *nup84Δ::Hyg* | Deletion of Nup84 using pFA6-hphNT1 | This study |
| yAV228 | yAV110; *nup53Δ::KanMX* | Deletion of Nup53 by PCR amplification of Nup53 deletion cassette from YKO collection | This study |
| yAV187 | BY *kap121-GFP::HIS3; pGal1-VHH[Nic96]-mKate2::URA* | Expression of VHH[Nic96]-mKate2 from a centromeric plasmid (pAV44) in a Kap121-GFP strain (GFP collection) | This study, ref GFP collection |
| yAV154 | yAV117; *atg15Δ::Hyg* | Deletion of *atg15* using pFA6-hphNT1 | This study |
| yAV164 | yAV121; *atg15Δ::Hyg* | Deletion of *atg15* using pFA6-hphNT1 | This study |
| yAV145 | *snf7Δ::Hyg; leu2::pGal1-VHH[Nup84]-mNeogreen::LEU2* | BWCPL1719 strain (gift from Lusk lab) expressing VHH[Nup84]-mNeogreen from an inducible galactose promoter, integrated in LEU locus using pAV49 | This study |
| yAV147 | *snf7Δ::Hyg; leu2::pGal1-VHH[Nic96]-mNeogreen::LEU2* | BWCPL1719 strain (gift from Lusk lab) expressing VHH[Nic96]-mNeogreen from an inducible galactose promoter, integrated in LEU locus using pAV50 | This study |
| Kap121-GFP | *BY4741 kap121-GFP::HIS3* | Genomic tagging of Kap121 | <sup>84</sup> |
| yAV264 | *yAV232; nup60-mTQ2::URA3* | C-terminal tagging of Nup60-mTQ2 using pSJ1458 | This study |
| yAV281 | *W303; brl1-AID::KanMX his3::osTIR1::HIS leu2::pGal1-VHH[Nic96]-mNeongreen::LEU2* | Genomic integration of VHH[Nic96]-mNeongreen under inducible galactose promoter using pAV50 in cells expressing brl1-AID::KanMX his3::osTIR1 (KWY9202; gift from Weis lab) | This study |
| KWY9269 | *W303; brl1-AID::KanMX his3::osTIR1::HIS trp1::ds RED-HDEL::TRP nup82-GFP::URA3* | Gift from Weis lab |  |
| yAV277 | *yAV110; nup157ΔC::NatNT2* | Deletion of Nup157 C-terminal (aa 628-1391) domain using pFA6-NatNT2 | This study |
| yAV278 | *yAV110; nup170ΔC::NatNT2* | Deletion of Nup170 C-terminal domain (aa 720 -1502) using pFA6-NatNT2 | This study |
| yAV266 | *diploid W303; leu2::pGal1-VHH[Nic96]-mNeongreen::LEU2* | Genomic integration of VHH[Nic96]-mNeongreen under inducible galactose promoter using pAV50 | This study |
| PMCPL1321 | *W303; leu2::pGal1-VHH[Nic96]-mNeongreen::LEU2 vph1-3xHA-mCherry::Nat* | Progeny of cross between yAV110 and PMCPL838. | This study |
| yAV227 | *W303; dga1Δ::TRP lro1Δ::HIS are1Δ::HIS are2Δ::HIS leu2::pGal1-VHH[Nic96]-mNG::LEU2* | YMG05 strain (gift from Lusk lab) expressing VHH[Nic96]-mNeongreen under inducible galactose promoter using pAV50 | This study |
| yAV280 | *BY pdr16-GFP::HIS3; pGal1-VHH[Nic96]-mKate2::URA* | Expression of VHH[Nic96]-mKate2 from a centromeric plasmid (pAV44) in a Pdr16-GFP strain (GFP collection) | This study, <sup>84</sup> |
| yAV286 | *BY erg6-GFP::HIS3; pGal1-VHH[Nic96]-mKate2::URA* | Expression of VHH[Nic96]-mKate2 from a centromeric plasmid (pAV44) in a Erg6-GFP strain (GFP collection) | This study, <sup>84</sup> |
| yAV287 | *BY ldo16-GFP::HIS3; pGal1-VHH[Nic96]-mKate2::URA* | Expression of VHH[Nic96]-mKate2 from a centromeric plasmid (pAV44) in a Ldo16-GFP strain (GFP collection) | This study, <sup>84</sup> |
| yAV290 | *yAV281; hmg1-mKate2::URA* | C-terminal tagging of Hmg1-mKate2 using pTO30 | This study |
| yAV295 | *yAV110; hmg1-mKate2::URA* | C-terminal tagging of Hmg1-mKate2 using pTO30 | This study |
| yAV296 | *yAV235; hmg1-mKate2::URA* | C-terminal tagging of Hmg1-mKate2 using pTO30 | This study |

**Table 2.** Plasmids.

| Name | Description | Source |
| --- | --- | --- |
| pRS303 | Integrative vector with HIS3 marker | Addgene <sup>85</sup> |
| pRS305 | Integrative vector with LEU2 marker | Addgene <sup>85</sup> |
| pAV49 | pGal1-VHH[Nup84]-mNeongreen-Cyc in pRS305 backbone | This study |
| pAV50 | pGal1-VHH[Nic96]-mNeongreen-Cyc in pRS305 backbone | This study |
| pAV70 | pNup120-VHH[Nic96]-mNeongreen-Cyc in pRS305 backbone | This study |
| pSJ1458 | pFA6-link-mTurquoise2-CaURA3MX | Sue Jaspersen (Addgene plasmid 86424) |
| pFA6-hphNT1 | Gene deletion cassette; hygromycin resistance | <sup>86</sup> |
| pFA6-natNT2 | Gene deletion cassette; clonNAT resistance | <sup>86</sup> |
| pPP014 | PCR template for C-terminal mCherry tagging; URA3 selection | <sup>42</sup> |
| pKW803 | pFA6-S-TEV-ZZ; HIS3 selection | This study |
| pAV61 | PCR template for C-terminal HALO; generated by replacing eYFP with a Halo-tag (pUC57) in pYM40 | This study |
| pAV64 | pGal1-VHH[Nic96]-HALO-Cyc in pRS305 backbone | This study |
| pAV69 | pGal1-VHH[Nic96]-mNeongreen-Cyc in pRS303 backbone | This study |
| pBT014 | Nab2NLS-mCherry; LEU selection | <sup>36</sup> |
| pAV54 | PCR template for C-terminal mNeongreen; generated by replacing mCherry with mNeongreen in pPP014 | <sup>57</sup> |
| pAV44 | Centromeric plasmid expressing pGal1-VHH[Nic96]-mKate2 generated by replacing pNup120-VHH[Nic96] with pGal1-VHH[Nic96] in TUS4991 | This study |
| pYM40 | PCR template for C-terminal YFP; hygromycin resistance | <sup>86</sup> |
| TUS4991 | Centromeric plasmid expressing pNup120-VHH[Nic96]-mKate2; URA selection | <sup>27</sup> |
| pAV79 | pGal1-Nic96-mNeongreen-Cyc in pRS305 backbone | This study |
| pAV80 | pGal1-Nup192-mNeongreen-Cyc in pRS305 backbone | This study |
| pAV81 | pGal1-Nup57-mNeongreen-Cyc in pRS305 backbone | This study |
| pTO30 | PCR template for C-terminal mKate2 tagging; URA3 selection | This study |

For imaging experiments, 2 days prior to the image acquisition (day-2), cells were grown at 30°C while shaking (200 rpm) in synthetic minimal medium supplemented with 2% D-glucose. For galactose induction experiments, cells were subsequently diluted in minimal medium supplemented with 2% D-raffinose on day-1 and grown to exponential phase. Cells were maintained in exponential growth phase from this point onwards. To induce VHH expression, cells were incubated for 20 min (unless indicated otherwise) with 0.5% D-galactose, followed by addition of 1% D-glucose. Expression of a Nab2-NLS reporter (Fig. 3A) was induced with 0.1% D-galactose for 3hrs. For constitutive expression experiments, cells were grown in 2% D-glucose for the entire experiment, maintaining exponential growth. Cells expressing centromeric plasmids were grown in appropriate selective minimal medium for the entire experiment. Cells in 1 ml of culture were concentrated to 50 µl by centrifugation (5000 rpm, 2 min) prior to imaging.

For spot assays, overnight cultures grown in YPD at 30°C were diluted to 0.2×10^7^ – 0.7×10^7^, recovered for 2-3 hours and subsequently diluted in a 10-fold dilution series to 0.5×10^6^ - 0.5×10^3^ in sterile water. 3 µl of serial dilutions were plated on YPD and YPGal plates and grown at 30°C for 1.5 days.

Brl1 depletion (Fig. 2E) was induced in exponentially growing cells (OD600 0.1-0.2) by addition of 0.5 uM auxin (indoleacetic acid; Sigma; I3750-5G-A, in ethanol) or the equivalent amount of ethanol to the medium. VHH[Nic96] expression was induced at t=3hrs as described above. Images were taken at t=4hr after the start of the experiment.

Growth rates were determined in synthetic dropout medium supplemented with 2% raffinose and 0.5% galactose. Cells were essentially grown as described under “strains and yeast growth” but grown to starvation overnight on day-1 to reach an OD600 of 3-4 at the beginning of the experiment. In the morning of day 0, cells were diluted to OD600 0.05 to start the experiment. Growth rates were determined by measuring OD600 every 1-1.5hrs.

### Replicative lifespan analysis

Replicative lifespan analysis was performed using microfluidic chips as previously described^39^. Cells were pre-cultured as described under “strains and yeast growth”. VHH[Nic96] expression was induced while loading the chip. Cells were maintained in synthetic dropout medium supplemented with 2% raffinose and 0.5% galactose for the entire experiment. Brightfield images (3 Z-slices of 0.7 µm apart) were taken every 20 minutes for a total time of 80 hours to follow cell division and replicative age. Fluorescent images of VHH[Nic96]-mNG expression were taken every 10 hours to ensure expression. Replicative lifespan data of BY4741 strains and BY4741 strains expressing GFP were taken from ^15^. To determine lifespan, only cells that could be followed their entire lifespan were included.

### Fluorescence microscopy

Images were acquired at 30°C using a Deltavision Elite (Applied Precision) microscope equipped with an Olympus UPlanSApo 100x (NA 1.4) oil immersion objective. An EDGE sCMOS5.5 camera was used for detection. Images were acquired in 30 Z-slices of 0.1 µm, except for Fig. 2C (3 z-slices of 0.5 µm). Timelapse imaging (video S1) was acquired by taking 3 successive Z-slices of 0.2 µm every 5 seconds for a total of 150 timepoints. All images were deconvolved using softWoRx software (GE Healthcare) and processed using Fiji software to generate images (ImageJ 2.14.0 ^66^).

AUTODOT^TM^ staining (Abcepta; SM1000a) was performed by addition of 1ul of dye to 1 ml of culture 15 minutes prior to imaging.

N/C ratios (Fig. 3A) were determined by measuring mean nuclear and cytosolic intensities in a defined area (10×10) of respectively nucleus and cytosol and correcting for background intensities. Cells that did not have mCherry signal twice above background were excluded from analysis.

Quantification of lipid droplet proximity (Fig. 6I) was performed by running PunctaFinder over each dataset (see below) and ranking VHH[Nic96]-mNG foci intensities in an descending order. The 50 highest-intensity foci were selected for manual inspection in Fiji and scored for lipid droplet proximity based on adjacent or overlapping signal in the AUTODOT^TM^ channel, either in the z-slice reported by PunctaFinder or in the z-slices above or below considering the full vertical range of each VHH[Nic96]-mNG focus. Proximity of VHH[Nic96]-mKate2 foci to GFP-tagged lipid droplet proteins (Fig. 6D) was quantified in a similar manner but with PunctaFinder settings optimized for mKate2 detection (see below). Cells that did not show signal in either of the two channels (VHH or AUTODOT^TM^/GFP) were excluded from analysis. In Brl1-depleted cells, NE localization (Fig. S6B) was determined by comparing VHH[Nic96] localization to Hmg1-mKate2 signal.

Cytosolic VHH[Nic96]-mNG intensities (Fig. S6A) were determined by measuring mean intensity in a defined cytosolic area (10×10) in sum projections of 30 z-slices and correcting for background intensities.

### STED microscopy

Cells were grown as described above. For live-cell labelling of yeast cells, cells were stained at the time of induction using 1 µM SiR-Halo dye for 1hr. Expression of VHH[Nic96]-Halo was shut off after 20 minutes by addition of 1% glucose to the medium, following a 40-minute recovery. 1 ml of culture was harvested (1 min, 4400 rpm) and cells were immediately used for live-cell imaging (stacked between a 18 mm ∅, borosilicate glass coverslip, No. 1.5H and a glass slide).

STED nanoscopy was performed at 30°C using a commercial microscope (Abberior Instruments GmbH), containing a STED laser (775 nm), four excitation lasers (640, 561, 448 and 405 nm) and equipped with a CoolLED pE-2 excitation system and a 100x oil immersion objective (Olympus UPLSAPO/1.40). STED images were taken using a pixel size of 18 nm, dwell time of 13 µs, 1.5% excitation using the 640 nm laser and a 660-730 nm detection window. 20% or 18% STED was applied for Nic96- or VHH-Halo strains respectively. Image acquisition was carried out using Imspector Software (v16.3, Abberior Instruments). Analysis was performed using Fiji Software (ImageJ 2.14.0 ^66^).

### CLEM

Correlative light and electron microscopy was performed as described in ^67^. Cells were cultured in the presence of 0.5% D-galactose for 20 minutes followed by the addition of 1% D-glucose as previously described. The cells were subsequently pelleted by centrifugation at ∼800 relative centrifugal force for 5 minutes and high-pressure frozen in the 200 µm indentation of an Al platelet (Engineering Office M. Wohlwend 241) using an HPM100 (Leica Microsystems). The samples were then freeze-substituted in 0.1% uranyl acetate and automated temperature control was used to finish the solution exchange and embedding in Lowicryl HM20 (Polysciences). A diamond knife (Diatome) was used to generate 250 µm thick sections collected on carbon-supported 200-mesh copper grids (Ted Pella 01840).

VHH[Nic96]-mNeongreen was visualized by fluorescence microscopy on a DeltaVision microscope (Applied Precision) equipped with a UPlanSapo ×100 1.4 numerical aperture oil immersion objective (Olympus) and an AURA light engine (Lumencor) and CoolSnapHQ charge-coupled device (Photometrics) camera. Protein A-coated gold beads (Cell Microscopy Core, University Medical Center Utrecht) were added to the grids to aid in the tilt-series alignment.

Tilt series were acquired using automated tilt-series acquisition via SerialEM 3.6.15 ^68^ on an electron microscope (FEI TF20) operating at 200kV from approximately −65° to +65° (1° increments) with a high-tilt tomography holder (Fishione Instruments 2020), a 100 µm objective aperture, a 150 µm C2 aperture and a 4,000 x 4,000 Eagle charge-coupled device (FEI) at a binned pixel size of 1.242. IMOD 4.11.24 ^69^ as used for automatic reconstruction ^70^, nonlinear anisotropic diffusion filtering and manual segmentation. The distance between perinuclear lipid droplets and the closest NPC was measured in IMOD ^69^.

### Affinity purifications and mass spectrometry

Affinity purifications and mass spectrometry analysis related to figure 1DE and Figure S1AB and Figure 1F and Figure S1CDE are separately described below.

### KARMA and proteomic data acquisition and analysis

Related to the affinity purification experiments presented in figure 1DE and Figure S1AB: KARMA experiments and data analysis were essentially performed as described in ^22^. To induce VHH bait expression, cells were grown to mid-log phase as described above and then switched from light L-lysine medium (25 mg/L) containing 2% raffinose to heavy, 13C6 15N2 L-lysine (Cambridge Isotope Laboratories) medium containing 2% galactose to concomitantly start VHH expression and metabolic labelling. After 20 minutes, bait expression was stopped by direct addition of 2% glucose (v/v) to the culture. Post-labelling samples were collected at the indicated timepoints by filtering the equivalent of 250 mL cells of OD600 of 1 and snap freezing them in liquid nitrogen.

All steps of the affinity purification protocol were carried out on ice, with minimal waiting times to preserve protein integrity. Frozen yeast pellets were resuspended in lysis buffer (20 mM HEPES pH 7.5, 50 mM KOAc, 20 mM NaCl, 2 mM MgCl2, 1 mM DTT, 10% v/v glycerol) and transferred to 2 mL screwcap microtubes (Sarstedt Inc.) preloaded with approximately 1 mL of 0.5 mm glass beads (BioSpec Products). The tubes were topped off with lysis buffer, and cells were lysed mechanically using a Mini BeadBeater-24 (BioSpec Products) in four 1-minute cycles at 3500 oscillations per minute, with 1-minute cooling intervals in ice water between cycles. Cell debris and unlysed cells were pelleted at 4°C for 30 s at 850 g and 1 mL of the remaining supernatant was supplemented with 110 µL 10× Detergent mix (protease inhibitor cocktail [Sigma-Aldrich], 5% v/v Triton x-100, 1% v/v Tween-20 in lysis buffer). This mixture was transferred to a fresh tube with 1 mg IgG-coated pre-equilibrated magnetic beads (Invitrogen Dynabeads M-270 Epoxy #14301; Sigma rabbit IgG serum #I5006). After incubating the affinity purification samples at 4°C for 30 minutes with continuous agitation, the beads were washed four times with 1 mL of wash buffer (0.1% v/v Tween-20 in lysis buffer). Proteins were then eluted in 40 µL of 1× Laemmli sample buffer for 2 minutes at 50°C. The eluates were subsequently denatured at 95°C for 5 minutes and snap-frozen in liquid nitrogen.

Eluted proteins were concentrated by SDS-PAGE using a 4% acrylamide stacking gel, then incubated in distilled water for 12 hours to remove residual detergents. Protein bands were excised and processed using a standard in-gel digestion protocol. Disulfide bonds were reduced with dithiothreitol (6.5 mM DTT in 100 mM ammonium bicarbonate) for 1 hour at 60°C, followed by alkylation with iodoacetamide (54 mM in 100 mM ammonium bicarbonate) for 30 minutes at 30°C in the dark. Proteins were then digested for 16 h at 37°C with 1.25 µg trypsin (Promega) in 100 mM ammonium bicarbonate. Peptides were purified using C18 BioPureSPN mini columns (The Nest Group, Inc), washed and desalted with Buffer A (0.1% v/v formic acid), eluted with Buffer B (0.1% v/v formic acid, 80% v/v acetonitrile), and finally recovered in 12.5 µL Buffer A containing iRT peptides (1:50 v/v, Biognosys).

LC-MS/MS analysis was performed on an Orbitrap Exploris 480 mass spectrometer (Thermo Scientific) coupled to a Vanquish Neo UHPLC system (Thermo Scientific). Peptides were separated using a C18 reversed phase column (75 μm x 400 mm (New Objective), packed in-house with ReproSil Gold 120 C18, 1.9 μm (Dr. Maisch GmbH)). For the 90-minute post-labelling time point, a 120-minute linear gradient from 7% to 35% buffer B (0.1% [v/v] formic acid, 80% [v/v] acetonitrile) was applied at a flow rate of 300 nl/min. The mass spectrometer was operated in data-independent acquisition (DIA) mode using one MS1 scan (350-1150 m/z, 120000 resolution, 250% normalized AGC target, 264 ms maximum injection time), followed by 41 variable MS2 windows from 350-1150 m/z with 1 m/z overlap (30000 resolution, 250% normalized AGC target, 64 ms maximum injection time). Fragmentation was performed by HCD at normalized collision energy (NCD) 28%. For the 180-minute post-labelling time point and label-free proteomic samples, data were acquired using a 60-minute non-linear gradient from 1% to 43.7% buffer B at a flow rate of 300 nl/min. DIA acquisition used one MS1 scan (330-1650 m/z, 120000 resolution, 300% normalized AGC target, 20 ms maximum injection time), followed by 30 variable MS2 windows from 330 to 1650 m/z with 1 m/z overlap (30000 resolution, 2000% normalized AGC target, 64 ms maximum injection time). Ions were fragmented with HCD (NCE 27%).

SILAC-DIA data from the 90-minute post-labelling time point was extracted with Spectronaut v18.2 (Biognosys) using a spectral library previously generated from 60 APs with 11 Nup baits ^23^. Data from the 180-minute post-labelling time point and the label-free DIA experiments were analysed with Spectronaut v19.1 using the directDIA workflow. For SILAC samples, the heavy channel (K+ 8.014) was generated in-silico using the “labelled” workflow. Carbamidomethylation was set as fixed modification and methionine oxidation, and N-terminal acetylation were included as variable modifications. Tryptic peptides with a maximum of two missed cleavages were considered and spectra were searched against the *Saccharomyces cerevisiae* protein database (downloaded from SGD on 13/10/2015, 6713 entries). For the data extraction in Spectronaut, default settings were used except that “cross-run normalization” was turned off. The ion intensities at the fragment level were exported and analysed in R as follows: First, low-quality precursors were excluded based on Spectronaut’s “F. ExcludedFromQuantification” flag. Only prototypic y-type fragment ions containing a single lysine residue and an intensity > 4 were retained. All remaining fragment ion intensities were summed separately for the heavy and light channel per precursor. Precursor-level fractional labelling was computed as Heavy / (Heavy + Light), and protein-level fractional labelling was obtained by taking the median fractional labelling of all precursors belonging to the same protein. Proteins with fewer than three identified precursor ions in all three replicates were excluded from downstream analysis. For label-free, samples the same filtering strategy was applied, except that both y- and b-type fragment ions in the light channel were used. Precursor intensities were summarized into protein intensities using the MaxLFQ algorithm ^71^. All samples were median-normalized, and the mean of three biological replicates was taken as the final protein intensity. Statistical significance was assessed using a two-tailed Student’s t-test.

### Affinity purifications of VHH[Nic96]-ZZ, Pom34-ZZ and proteomic data acquisition and analysis

Related to the affinity purification experiments presented in Figure 1F and Figure S1CDE: affinity purification experiments of Pom34-ZZ and no ZZ (Fig. S1E, 6 replicates each) and VHH[Nic96]-ZZ (Fig. 1F, Fig. S1CD, 3 replicates) were essentially performed as in ^12^. In short, 3 liter cultures of strains yAV235 (no ZZ), yAV255 or KWY12172 were grown to an OD600 of ± 1.0 in YPD, pelleted (5000g; 5min at 4°C) and washed twice in ice-cold water (2000g; 5min at 4°C) and once in buffer (20 mM K-HEPES (pH 7.4), 1.2% PVP, 1mM DTT, supplemented with 1:200 solution P (2mg PepstatinA + 90 mg PMSF in 5 ml EtOH) and protease inhibitor cocktail (EDTA-free protease inhibitor cocktail cOmplete, Roche; 04693159001). Cells were centrifuged at 2500g for 20min at 4°C and the concentrated cell slurry was snap frozen in liquid nitrogen and stored at −80°C. Cells were pulverized into fine powder via cryogenic milling in a Retch planetary ball mill (PM100) as described in ^72^; frozen powder was stored at −80°C.

For affinity purifications, 0.5g of frozen cell powder was resuspended in 3 ml RT extraction buffer (20 mM HEPES, 110 mM KAc, 2 mM MgCl2, 0.1% Triton-×100, 0.1% Tween-20, 150 mM NaCl) supplemented with 1:200 solution P, protease inhibitor cocktail and 1mM DTT. The suspension was homogenized three times with a Polytron (Kinematica AG) at 5.5 setting for 30 seconds, cooling the suspension 30 seconds on ice in between sessions. Cell lysates were isolated by centrifugation (2000g; 15 min at 4°C) and added to IgG-coated pre-equilibrated magnetic beads (Invitrogen Dynabeads M-270 Epoxy #14301; Sigma rabbit IgG serum #I5006) as described in ^73^. Samples were incubated for 1hr at 4°C with end-over-end mixing. To elute, beads were washed three times with ice-cold buffer (20 mM HEPES, 110 mM KAc, 2 mM MgCl2, 0.1% Triton-×100, 0.1% Tween-20, 150 mM NaCl) and incubated for 10min with 2% SDS in 40mM Tris pH 7.4 at 50°C. Elution fractions were subsequently analysed by SDS-page, immunoblot or proteomics as indicated.

Affinity purifications of VHH[Nic96]-ZZ (Fig. 1F, Fig. S1CD) were essentially performed as above, using the corresponding buffers as described in the legend of Fig. S1C and in smaller volumes (200mg frozen cell powder in 800uL extraction buffer) and omitting the Polytron homogenizing step. Elution fractions of affinity purifications using buffer 7 (1.5M Ammonium Acetate pH 7.0, 1% Triton X-100, supplemented with protease inhibitor cocktail, solution P and 1mM DTT) were used for subsequent analysis (Fig. 1F, Fig. S1D).

To prepare samples for proteomic analysis, DTT (50 mM final concentration) was added to elution fractions from affinity purification experiments and samples were loaded into 4-12% Bis-Tris gels (Invitrogen^TM^) with MOPS buffer. Samples were run for 10 min at 200V and stained in Coomassie blue overnight. Gel plugs were then destained and sliced into small pieces, washed in 50% v/v acetonitrile in 100mM ammonium bicarbonate and incubated for 30 minutes, followed by a 5min wash with 100% v/v acetonitrile. Gel pieces were subsequently dried at 37°C. Disulfide bonds were reduced with 10 mM DTT and proteins were alkylated with 55 mM iodoacetamide, washed once more for 5min with 100% v/v acetonitrile followed by overnight digestion in trypsin (sequencing grade modified trypsin V5111, Promega) at 37°C. Peptides were eluted with 75% v/v acetonitrile plus 5% v/v formic acid (20 min; 500 rpm). Eluted peptides were dried under vacuum and resuspended in 0.1% (v/v) formic acid.

Discovery mass spectrometric analysis was performed on a quadruple orbital mass spectrometer equipped with a nano-electrosrpay ion source (Orbitrap Exploris 480, Thermo Scientific). Peptides were separated by liquid chromatography on a Evosep system (Evosep One, Evosep) using a nano-LC column. LC-MS raw data were processed with Spectronaut (version 19.0.240624; Biognosys) with standard settings of the directDIA workflow.

Data analysis for enrichment in Pom34-ZZ APs was performed using the Bioconductor DEP package^74^ in R. We used a stringent filtering criterion to exclude proteins with missing values in all 6 replicas of either of the two conditions from analysis. Missing values were subsequently imputed using the MinProb imputation method to calculate log2-fold enrichment in the Pom34-ZZ purifications compared to control.

### Cell lysates for immunoblotting

For determining VHH expression levels (Fig. S3B), 20 ml of culture of ∼OD600 1.0 were harvested at t_on_=90min. Cells were pelleted at 4000 rpm for 5 minutes at 4°C, washed once with PBS and frozen in liquid nitrogen. For lysis, the cell pellet was resuspended in HEPES buffer (20 mM HEPES, 110 mM KAc, 2 mM MgCl2, 0.1% Triton-×100, 0.1% Tween-20, 150 mM NaCl) supplemented with protease inhibitor cocktail (EDTA-free protease inhibitor cocktail cOmplete, Roche; 04693159001) and transferred to 1.5 ml screw cups containing glass beads (0.5mm). Samples were lysed using a FastPrep following standard yeast protocols (FastPrep MP Biomedicals), with 5-minute cooling on ice in between. Glass beads and cell debris were pelleted by centrifugation (6000 rpm; 5 min at 4C) and the cell lysate was subsequently centrifuged for 15 min at 13000 rpm at 4 °C. Protein concentration was determined using a Pierce BCA protein assay kit (Thermo Scientific^TM^, #23235).

### Immunoblotting

Cell lysates were diluted in 4x LDS sample buffer containing 50mM DTT, heated for 10 min at 70°C, centrifuged (15 000 rpm; 2 min) and separated by SDS-page (NuPAGE^TM^ 4-12% Bis-Tris Mini Protein Gels, Invitrogen^TM^). SDS-page and transfers were done according to standard immunoblotting protocols, using a PVDF membrane. Membranes were blocked for 1-2 hour at RT with 5% milk (w/v%) in TBS-Tween (v/v 0.1%) (TBST), then incubated overnight at 4C with primary antibody listed below. Membranes were washed three times for 10 min with TBST and subsequently incubated for 30-60 minutes with secondary antibodies at 4C. Membranes were washed three times for 10min with TBST and imaged using a ECL development system (Millipore WBLUF0500). The following antibodies were used: 1:2,000 Rabbit Isotype Control DA1E (Cell Signaling CST3900S); VHH-ZZ detection (Fig. S3B)), 1:10,000 HRP-linked anti-rabbit (Cytiva, NA934V). All antibody dilutions were made in 5% BSA in TBST, 0.02% w/v sodium azide.

### PunctaFinder analysis

Quantification of VHH[Nic96]-mNG intensities and Nic96-mNG intensities were performed using the PunctaFinder algorithm as described in ^44^. In brief, cell masks were generated using the ImageJ plugin BudJ ^75^. Cells that did not show detectable mNG expression, daughter cells without a visible NE, dividing cells with a mitotic bridge and dead cells were all excluded from analysis. We also excluded cells with the midplane of the NE in the first 5 or last 5 z-slices and cells near the edge of the image. Following visual inspection of VHH[Nic96]-mNG foci, puncta diameter was set at 5 pixels (1/15.4943 um/pixel). To determine threshold values for VHH[Nic96]-mNeongreen detection, we created a validation dataset of 91 images (sampling timepoint t=2hr; 20-minute induction pulse as described in Fig. 2A) containing 453 puncta and performed 5 bootstrap iterations (with a weighting factor for false negatives of 0.75) to determine optimal threshold values. The following threshold values were used for VHH[Nic96]-mNG foci detection: T_local_ = 1.478 (ratio punctum/surroundings) and T_global_ = 1.816 (ratio punctum/cell)). T_cv_ was omitted as it prevented detection of the brightest VHH[Nic96] foci in the NE.

After establishing optimal thresholds, we then run PunctaFinder over the full dataset, selecting cells based on the specified criteria. Z-slices 1-3 and 28-30 were excluded from analysis to exclude deconvolution artifacts. For optimal foci detection in *xyz* dimensions, candidate puncta were selected based on weighted T_local_ (T_local_ + 0.15 * T_global_) in *xy* and T_global_ in *z*, allowing minimal overlap (1 pixel both in *xy* and *z*) between puncta. We visually validated the detection of at least 350 VHH[Nic96]-mNG foci (both constitutively expressed and galactose induction) in >10 cells. PunctaFinder output was analysed in R. To further reduce the number of false positives, we manually excluded puncta that were not ∼3x above background (intensity < 500). Nic96-mNG (Fig. 5D) was performed with similar PunctaFinder settings but subsequent punctum filtering was omitted, as the absence of cytosolic signal did not require additional filtering steps.

For detection of VHH[Nic96]-mKate2 foci (Fig. 6D), we used T_local_ = 1.37 and T_global_ = 1.57 without additional filtering steps. All the detected foci were manually validated as they were scored for LD proximity.

### Modelling of VHH[Nic96] labelling over time

We used generalized linear mixed models (GLMMs) to understand the relationship between nr of VHH[Nic96]-mNG foci and time following the short induction pulse in two strain backgrounds (haploid versus diploid cells)

To model the number of foci in time, we fitted a negative-binomial GLMM with experimental replica number (four for each of the two strains) included as a random effect to predict the number of foci per cell over time. We fitted the number of VHH[Nic96] foci as a polynomial with regards to time up to the second order. We additionally examined the fit under a Poisson distribution using the DHARMa package ^76,77^, but that resulted in overdispersion.

All models were fitted using the glmmTMB package in R ^78^. The interaction between time and strain and between time squared and strain were both significant (ANOVA, Wald chisquare tests; Supplementary table 1). To validate the fitted models, we checked the residuals with the DHARMa package^76^ to check for overdispersion and zero inflation (see Fig. S5 DEF).

Model formulas for the selected model (haploid and diploid): for the i^th^ observation of the j^th^ replica, the number of foci within that cell comes from a negative binomial distribution with mean *μ* and shape parameter *θ*, following the nbinom2 family as implemented in glmmTMB:

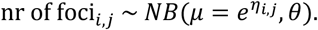

The value of the linear predictor of the mean is given by:

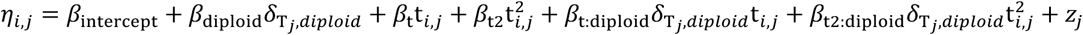

here, the β coefficients are estimated during the model fitting, t*_i_*_,*j*_ is the time for the i^th^ observation of the j^th^ replica, *δ*_T*j*,*diploid*_ is 0 if the strain of the j^th^ replicate is haploid and 1 if the strain is diploid, and *z_j_* corresponds to a contribution from a random effect at the replicate level, which is drawn from a normal distribution with standard deviation *σ_replica_*:

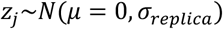

The coefficient estimates and standard errors (when estimated) are shown in Supplementary Table 2. The confidence interval of Δt_max,haploid - diploid_ was approximated by calculating the difference for 10^5^ sampled sets of coefficients, each of which was drawn from a multivariate normal distribution with the coefficient values as mean and variance covariance set to the variance covariance matrix of the parameters from the fit.

### Structural model

To check for potential structural clashes of the VHH[Nic96] in the NPC, the X-ray structure of Nic96 bound by VHH[Nic96] (PDB: 6X07, ^27^) was superimposed with the recent high-resolution cryo-EM structure of the inner spoke ring of the yeast NPC (PDB: 8TJ5, ^28^ and 7N9F, ^10^) using the matchmaker command in ChimeraX.

### Statistics

Biological and technical replicates are indicated in figure legends. Graphs and statistical analysis were generated using R version 4.1.0, 4.4.0 and 4.5.2 ^79^. Depending on the response variable, we applied different modelling approaches. For count data (Fig. 2H), a negative binomial GLMM was fitted using the glmmTMB package ^78^, including strain background as predictors and replica as random effect. For proportional data (Fig. 6I, Fig. S6C), we fitted binomial GLMs, including treatment (Fig. 6I) or NE localization (Fig. S6C) as predictor and replica as random effect. For foci intensities (Fig. 2I), we fitted a linear mixed-effects model using the lme4 package ^80^, including strain background as a predictor and replica as random effect. We applied a log transformation to the response variable to improve model assumptions. In all cases, model assumptions were checked using diagnostic plots from the DHARMa package ^76^.

ANOVA tables were obtained using the car package ^81^ or lmerTest ^82^. Post hoc comparisons (Fig. 6I) were performed using estimated marginal means (emmeans) ^83^, with pairwise comparisons (trt.vs.ctrl; ctrl = WT (pulsed) or exponential (constitutive)) and Dunnett-adjusted p-values to account for multiple testing.

## Data availability

The STED and CLEM imaging data and mass spectrometry data that support the findings in this manuscript will be deposited in public databases. Source data to the figures are provided within the paper or are available upon request.

## Acknowledgements

We kindly thank Prof. Dr. Thomas Schwartz (MIT) for sharing the Nic96 and Nup84 nanobodies. We kindly thank Dr. Alexey N. Butkevich and Prof. Dr. Stefan W. Hell (Max Planck Institute for Medical Research, Heidelberg) for providing the SiR-Halo dye. We are grateful to Nyassa de Kock Jewell, Kyra Belder, Karin Wolters, and Mariska de Graaff for their invaluable support with the affinity purifications and mass spectrometry. We thank Federico Uliana and Ino D Karemaker from the IBC MS facility. We thank Michael Chang and all members of the Chang and Veenhoff labs for valuable input. Annemiek Veldsink was supported by a PhD-fellowship from the Graduate School of Medical Sciences of the University of Groningen. This work was supported by a Vici grant (VI.C.192.031) to LMV from the Netherlands Organisation for Scientific Research and funding to KW from the Swiss National Science Foundation (CRSII5_193740).

## Author contribution

ACV and LMV conceived and designed the project with input from all authors. ACV performed and analysed all experiments alone or in collaboration with the co-authors. Specifically, ACV, JSF and SH performed the KARMA experiments supervised by KW; ACV, HMT, and MH optimized the use of Puncta Finder and KJvB provided expertise in statistical modelling; ACV, EMFdL performed STED supervised by RV, and ACV and LJS performed affinity purifications. The CLEM experiments were performed by PJM supervised by PCL. ACV, and LMV wrote the manuscript with input from all authors.

## Materials and Correspondence

Materials and correspondence may be addressed to Liesbeth Veenhoff

## Supplementary Information

Figure S1-S6

Table S1-S2

Supplementary video S1

## References

1 Petrovic, S., Mobbs, G. W. & Hoelz, A. Structure, function and assembly of nuclear pore complexes. Nature Reviews Molecular Cell Biology 27, 35–54 (2026). 10.1038/s41580-025-00881-w

2 Cowburn, D. & Rout, M. Improving the hole picture: towards a consensus on the mechanism of nuclear transport. Biochem Soc Trans 51, 871–886 (2023). 10.1042/bst20220494

3 Wing, C. E., Fung, H. Y. J. & Chook, Y. M. Karyopherin-mediated nucleocytoplasmic transport. Nature Reviews Molecular Cell Biology 23, 307–328 (2022). 10.1038/s41580-021-00446-7

4 Simon, M. N., Dubrana, K. & Palancade, B. On the edge: how nuclear pore complexes rule genome stability. Curr Opin Genet Dev 84, 102150 (2024). 10.1016/j.gde.2023.102150

5 Sakuma, S. et al. Inhibition of Nuclear Pore Complex Formation Selectively Induces Cancer Cell Death. Cancer Discovery 11, 176–193 (2021). 10.1158/2159-8290.Cd-20-0581

6 Coyne, A. N. et al. Nuclear accumulation of CHMP7 initiates nuclear pore complex injury and subsequent TDP-43 dysfunction in sporadic and familial ALS. Sci Transl Med 13 (2021). 10.1126/scitranslmed.abe1923

7 Kim, S. J. et al. Integrative structure and functional anatomy of a nuclear pore complex. Nature 555, 475–482 (2018). 10.1038/nature26003

8 Petrovic, S. et al. Architecture of the linker-scaffold in the nuclear pore. Science 376, eabm9798 (2022). doi:10.1126/science.abm9798

9 Zimmerli, C. E. et al. Nuclear pores dilate and constrict in cellulo. Science 374, eabd9776 (2021). doi:10.1126/science.abd9776

10 Akey, C. W. et al. Comprehensive structure and functional adaptations of the yeast nuclear pore complex. Cell 185, 361–378 e325 (2022). 10.1016/j.cell.2021.12.015

11 Schrader, N. et al. Structural basis of the nic96 subcomplex organization in the nuclear pore channel. Mol Cell 29, 46–55 (2008). 10.1016/j.molcel.2007.10.022

12 Alber, F. et al. The molecular architecture of the nuclear pore complex. Nature 450, 695–701 (2007). 10.1038/nature06405

13 Winey, M., Yarar, D., Giddings, T. H., Jr. & Mastronarde, D. N. Nuclear pore complex number and distribution throughout the Saccharomyces cerevisiae cell cycle by three-dimensional reconstruction from electron micrographs of nuclear envelopes. Mol Biol Cell 8, 2119–2132 (1997). 10.1091/mbc.8.11.2119

14 Makio, T. et al. The nucleoporins Nup170p and Nup157p are essential for nuclear pore complex assembly. Journal of Cell Biology 185, 459–473 (2009). 10.1083/jcb.200810029

15 Hurt, E. C. A novel nucleoskeletal-like protein located at the nuclear periphery is required for the life cycle of Saccharomyces cerevisiae. Embo j 7, 4323–4334 (1988). 10.1002/j.1460-2075.1988.tb03331.x

16 Lautier, O. et al. Co-translational assembly and localized translation of nucleoporins in nuclear pore complex biogenesis. Mol Cell (2021). 10.1016/j.molcel.2021.03.030

17 Seidel, M. et al. Co-translational assembly orchestrates competing biogenesis pathways. Nature Communications 13, 1224 (2022). 10.1038/s41467-022-28878-5

18 Kuiper, E. F. E. et al. The chaperone DNAJB6 surveils FG-nucleoporins and is required for interphase nuclear pore complex biogenesis. Nat Cell Biol 24, 1584–1594 (2022). 10.1038/s41556-022-01010-x

19 Prophet, S. M. et al. Atypical nuclear envelope condensates linked to neurological disorders reveal nucleoporin-directed chaperone activities. Nat Cell Biol 24, 1630–1641 (2022). 10.1038/s41556-022-01001-y

20 Verzijlbergen, K. F. et al. Recombination-induced tag exchange to track old and new proteins. Proc Natl Acad Sci U S A 107, 64–68 (2010). 10.1073/pnas.0911164107

21 Zsok, J. et al. Nuclear basket proteins regulate the distribution and mobility of nuclear pore complexes in budding yeast. Mol Biol Cell 35, ar143 (2024). 10.1091/mbc.E24-08-0371

22 Onischenko, E. et al. Maturation Kinetics of a Multiprotein Complex Revealed by Metabolic Labeling. Cell 183, 1785–1800.e1726 (2020). 10.1016/j.cell.2020.11.001

23 Kralt, A. et al. An amphipathic helix in Brl1 is required for nuclear pore complex biogenesis in S. cerevisiae. Elife 11 (2022). 10.7554/eLife.78385

24 Solà Colom, M., et al. A checkpoint function for Nup98 in nuclear pore formation suggested by novel inhibitory nanobodies. Embo j 43, 2198–2232 (2024). 10.1038/s44318-024-00081-w

25 Otsuka, S. et al. A quantitative map of nuclear pore assembly reveals two distinct mechanisms. Nature 613, 575–581 (2023). 10.1038/s41586-022-05528-w

26 Latham, A. P. et al. Integrative spatiotemporal modeling of biomolecular processes: Application to the assembly of the nuclear pore complex. Proceedings of the National Academy of Sciences 122, e2415674122 (2025). doi:10.1073/pnas.2415674122

27 Nordeen, S. A. et al. A nanobody suite for yeast scaffold nucleoporins provides details of the nuclear pore complex structure. Nat Commun 11, 6179 (2020). 10.1038/s41467-020-19884-6

28 Akey, C. W. et al. Implications of a multiscale structure of the yeast nuclear pore complex. Molecular Cell 83, 3283–3302.e3285 (2023). 10.1016/j.molcel.2023.08.025

29 Veldsink, A. C., Fischer, J. S., Hell, S., Weis, K. & Veenhoff, L. M. A tool to pulse-label yeast Nuclear Pore Complexes in imaging and biochemical experiments. (2025). 10.7554/elife.108399.1

30 Kosova, B., Panté, N., Rollenhagen, C. & Hurt, E. Nup192p Is a Conserved Nucleoporin with a Preferential Location at the Inner Site of the Nuclear Membrane *. Journal of Biological Chemistry 274, 22646–22651 (1999). 10.1074/jbc.274.32.22646

31 Hakhverdyan, Z. et al. Rapid, optimized interactomic screening. Nature Methods 12, 553–560 (2015). 10.1038/nmeth.3395

32 Guerra, P., Vuillemenot, L.-A., Rae, B., Ladyhina, V. & Milias-Argeitis, A. Systematic In Vivo Characterization of Fluorescent Protein Maturation in Budding Yeast. ACS Synthetic Biology 11, 1129–1141 (2022). 10.1021/acssynbio.1c00387

33 Doye, V., Wepf, R. & Hurt, E. C. A novel nuclear pore protein Nup133p with distinct roles in poly(A)+ RNA transport and nuclear pore distribution. Embo j 13, 6062–6075 (1994). 10.1002/j.1460-2075.1994.tb06953.x

34 Fernandez-Martinez, J. et al. Structure-function mapping of a heptameric module in the nuclear pore complex. J Cell Biol 196, 419–434 (2012). 10.1083/jcb.201109008

35 Fischer, J. S. et al. A conserved mechanism of membrane fusion in nuclear pore complex assembly. bioRxiv, 2025.2007.2021.665908 (2025). 10.1101/2025.07.21.665908

36 Timney, B. L. et al. Simple kinetic relationships and nonspecific competition govern nuclear import rates in vivo. J Cell Biol 175, 579–593 (2006). 10.1083/jcb.200608141

37 Hodge, C. A. et al. Integral membrane proteins Brr6 and Apq12 link assembly of the nuclear pore complex to lipid homeostasis in the endoplasmic reticulum. Journal of Cell Science 123, 141–151 (2010). 10.1242/jcs.055046

38 Onischenko, E. et al. Natively Unfolded FG Repeats Stabilize the Structure of the Nuclear Pore Complex. Cell 171, 904–917.e919 (2017). 10.1016/j.cell.2017.09.033

39 Scarcelli, J. J., Hodge, C. A. & Cole, C. N. The yeast integral membrane protein Apq12 potentially links membrane dynamics to assembly of nuclear pore complexes. Journal of Cell Biology 178, 799–812 (2007). 10.1083/jcb.200702120

40 Lord, C. L. & Wente, S. R. Nuclear envelope-vacuole contacts mitigate nuclear pore complex assembly stress. J Cell Biol 219 (2020). 10.1083/jcb.202001165

41 Webster, B. M., Colombi, P., Jager, J. & Lusk, C. P. Surveillance of nuclear pore complex assembly by ESCRT-III/Vps4. Cell 159, 388–401 (2014). 10.1016/j.cell.2014.09.012

42 Rempel, I. L. et al. Age-dependent deterioration of nuclear pore assembly in mitotic cells decreases transport dynamics. eLife 8, 1–26 (2019). 10.7554/eLife.48186

43 Lusk, C. P., Makhnevych, T., Marelli, M., Aitchison, J. D. & Wozniak, R. W. Karyopherins in nuclear pore biogenesis: a role for Kap121p in the assembly of Nup53p into nuclear pore complexes. Journal of Cell Biology 159, 267–278 (2002). 10.1083/jcb.200203079

44 Terpstra, H. M. et al. PunctaFinder: An algorithm for automated spot detection in fluorescence microscopy images. Mol Biol Cell 35, mr9 (2024). 10.1091/mbc.E24-06-0254

45 Jorgensen, P. et al. The size of the nucleus increases as yeast cells grow. Mol Biol Cell 18, 3523–3532 (2007). 10.1091/mbc.e06-10-0973

46 de Godoy, L. M. et al. Comprehensive mass-spectrometry-based proteome quantification of haploid versus diploid yeast. Nature 455, 1251–1254 (2008). 10.1038/nature07341

47 Klis, F. M., de Koster, C. G. & Brul, S. Cell wall-related bionumbers and bioestimates of Saccharomyces cerevisiae and Candida albicans. Eukaryot Cell 13, 2–9 (2014). 10.1128/EC.00250-13

48 Eisenberg-Bord, M. et al. Identification of seipin-linked factors that act as determinants of a lipid droplet subpopulation. Journal of Cell Biology 217, 269–282 (2017). 10.1083/jcb.201704122

49 Sandager, L. et al. Storage Lipid Synthesis Is Non-essential in Yeast *. Journal of Biological Chemistry 277, 6478–6482 (2002). 10.1074/jbc.M109109200

50 Kumanski, S., Viart, B. T., Kossida, S. & Moriel-Carretero, M. Lipid Droplets Are a Physiological Nucleoporin Reservoir. Cells 10 (2021). 10.3390/cells10020472

51 Seidel, M. et al. Co-translational binding of importins to nascent proteins. Nature Communications 14, 3418 (2023). 10.1038/s41467-023-39150-9

52 Diep, D. T. V. et al. A metabolically controlled contact site between vacuoles and lipid droplets in yeast. Developmental Cell 59, 740–758.e710 (2024). 10.1016/j.devcel.2024.01.016

53 Ren, J. et al. A phosphatidylinositol transfer protein integrates phosphoinositide signaling with lipid droplet metabolism to regulate a developmental program of nutrient stress–induced membrane biogenesis. Molecular Biology of the Cell 25, 712–727 (2014). 10.1091/mbc.e13-11-0634

54 Petschnigg, J. et al. Good fat, essential cellular requirements for triacylglycerol synthesis to maintain membrane homeostasis in yeast. J Biol Chem 284, 30981–30993 (2009). 10.1074/jbc.M109.024752

55 Garcia, E. J. et al. Membrane dynamics and protein targets of lipid droplet microautophagy during ER stress-induced proteostasis in the budding yeast, Saccharomyces cerevisiae. Autophagy 17, 2363–2383 (2021). 10.1080/15548627.2020.1826691

56 Hamed, M. & Antonin, W. Dunking into the Lipid Bilayer: How Direct Membrane Binding of Nucleoporins Can Contribute to Nuclear Pore Complex Structure and Assembly. Cells 10 (2021). 10.3390/cells10123601

57 Otto, T. A. et al. Nucleoporin Nsp1 surveils the phase state of FG-Nups. Cell Rep 43, 114793 (2024). 10.1016/j.celrep.2024.114793

58 Milles, S. et al. Facilitated aggregation of FG nucleoporins under molecular crowding conditions. EMBO Rep 14, 178–183 (2013). 10.1038/embor.2012.204

59 Rouviere, J. O. et al. A SUMO-dependent feedback loop senses and controls the biogenesis of nuclear pore subunits. Nat Commun 9, 1665 (2018). 10.1038/s41467-018-03673-3

60 Sakuma, S. et al. Homeostatic regulation of nucleoporins is a central driver of nuclear pore biogenesis. Cell Rep 44, 115468 (2025). 10.1016/j.celrep.2025.115468

61 Lone, M. A. et al. Yeast Integral Membrane Proteins Apq12, Brl1, and Brr6 Form a Complex Important for Regulation of Membrane Homeostasis and Nuclear Pore Complex Biogenesis. Eukaryot Cell 14, 1217–1227 (2015). 10.1128/ec.00101-15

62 Maslennikova, D. et al. Dystonia-associated Torsins sustain CLCC1 function to promote membrane fusion of the nuclear envelope for NPC biogenesis. bioRxiv, 2025.2011.2007.687155 (2025). 10.1101/2025.11.07.687155

63 Mathiowetz, A. J. et al. CLCC1 promotes hepatic neutral lipid flux and nuclear pore complex assembly. Nature (2026). 10.1038/s41586-025-10064-4

64 Diep, D. T. V. & Bohnert, M. The Vacuole Lipid Droplet Contact Site vCLIP. Contact (Thousand Oaks) 7, 25152564241308722 (2024). 10.1177/25152564241308722

65 Kohler, V. & Büttner, S. Remodelling of Nucleus-Vacuole Junctions During Metabolic and Proteostatic Stress. Contact 4 (2021). 10.1177/25152564211016608

66. Schindelin, J., et al. Fiji: an open-source platform for biological-image analysis. Nature Methods 9, 676–682 (2012). 10.1038/nmeth.2019

67 Mannino, P. J. et al. A quantitative ultrastructural timeline of nuclear autophagy reveals a role for dynamin-like protein 1 at the nuclear envelope. Nat Cell Biol 27, 464–476 (2025). 10.1038/s41556-025-01612-1

68 Mastronarde, D. N. Automated electron microscope tomography using robust prediction of specimen movements. J Struct Biol 152, 36–51 (2005). 10.1016/j.jsb.2005.07.007

69 Kremer JR, M. D., McIntosh JR. Computer visualization of three-dimensional image data using IMOD. J Struct Biol. 116, 71–76 doi:10.1006/jsbi.1996.0013

70 Mastronarde, D. N. & Held, S. R. Automated tilt series alignment and tomographic reconstruction in IMOD. J Struct Biol 197, 102–113 (2017). 10.1016/j.jsb.2016.07.011

71 Cox, J. et al. Accurate proteome-wide label-free quantification by delayed normalization and maximal peptide ratio extraction, termed MaxLFQ. Mol Cell Proteomics 13, 2513–2526 (2014). 10.1074/mcp.M113.031591

72 Cryogenic Disruption of Yeast Cells according to the Rout Protocol, <https://www.retsch.com/files/9985/cryogenic-disruption-of-yeast-cells-according-to-the-rout-protocol.pdf>

73 Conjugation of Dynabeads with Rabbit IgG, <https://commonfund.nih.gov/sites/default/files/Conjugation-of-Dynabeads.pdf>

74 Zhang, X. et al. Proteome-wide identification of ubiquitin interactions using UbIA-MS. Nature Protocols 13, 530–550 (2018). 10.1038/nprot.2017.147

75 Ferrezuelo, F. et al. The critical size is set at a single-cell level by growth rate to attain homeostasis and adaptation. Nature communications 3, 1012 (2012).

76 Hartig, Florian; DHARMa: Residual Diagnostics for Hierarchical (Multi-Level / Mixed) Regression Models.

77. Zuur, A., Ieno, E., Walker, N., Saveliev, A. & Smith, G. Mixed Effect Models and Extensions in Ecology With R. (2009).

78 Brooks, M. E. et al. glmmTMB Balances Speed and Flexibility Among Packages for Zero-inflated Generalized Linear Mixed Modeling. The R Journal 9, 378–400 (2017).

79 R Core Team; R: A Language and Environment for Statistical Computing (R Foundation for Statistical Computing, Vienna, Austria, 2024).

80 Bates, D., Mächler, M., Bolker, B. & Walker, S. Fitting Linear Mixed-Effects Models Using lme4. Journal of Statistical Software 67, 1 – 48 (2015). 10.18637/jss.v067.i01

81 Fox, J. & Weisberg, S. An R Companion to Applied Regression. Third edn, (Sage, 2019).

82 Kuznetsova, A., Brockhoff, P. B. & Christensen, R. H. B. lmerTest Package: Tests in Linear Mixed Effects Models. Journal of Statistical Software 82, 1 – 26 (2017). 10.18637/jss.v082.i13

83. emmeans: Estimated Marginal Means, aka Least-Squares Means v. R package version 2.0.3 (2026).

84 Huh, W. K., Falvo, J. V., Gerke, L. C., Carroll, A. S., Howson, R. W., Weissman, J. S., & O’Shea, E. K. Global analysis of protein localization in budding yeast. Nature 425, 686–691 doi:10.1038/nature02026

85 Markus, S. M., Punch, J. J. & Lee, W. L. Motor- and tail-dependent targeting of dynein to microtubule plus ends and the cell cortex. Curr Biol 19, 196–205 (2009). 10.1016/j.cub.2008.12.047

86 Janke, C. et al. A versatile toolbox for PCR-based tagging of yeast genes: new fluorescent proteins, more markers and promoter substitution cassettes. Yeast 21, 947–962 (2004). 10.1002/yea.1142

