## Supplement for "Detecting Nuclear Pore Complex assembly in living cells"

**Supplementary data to Veldsink et al**

**Figure S1-S6**

**Table S1, S2**

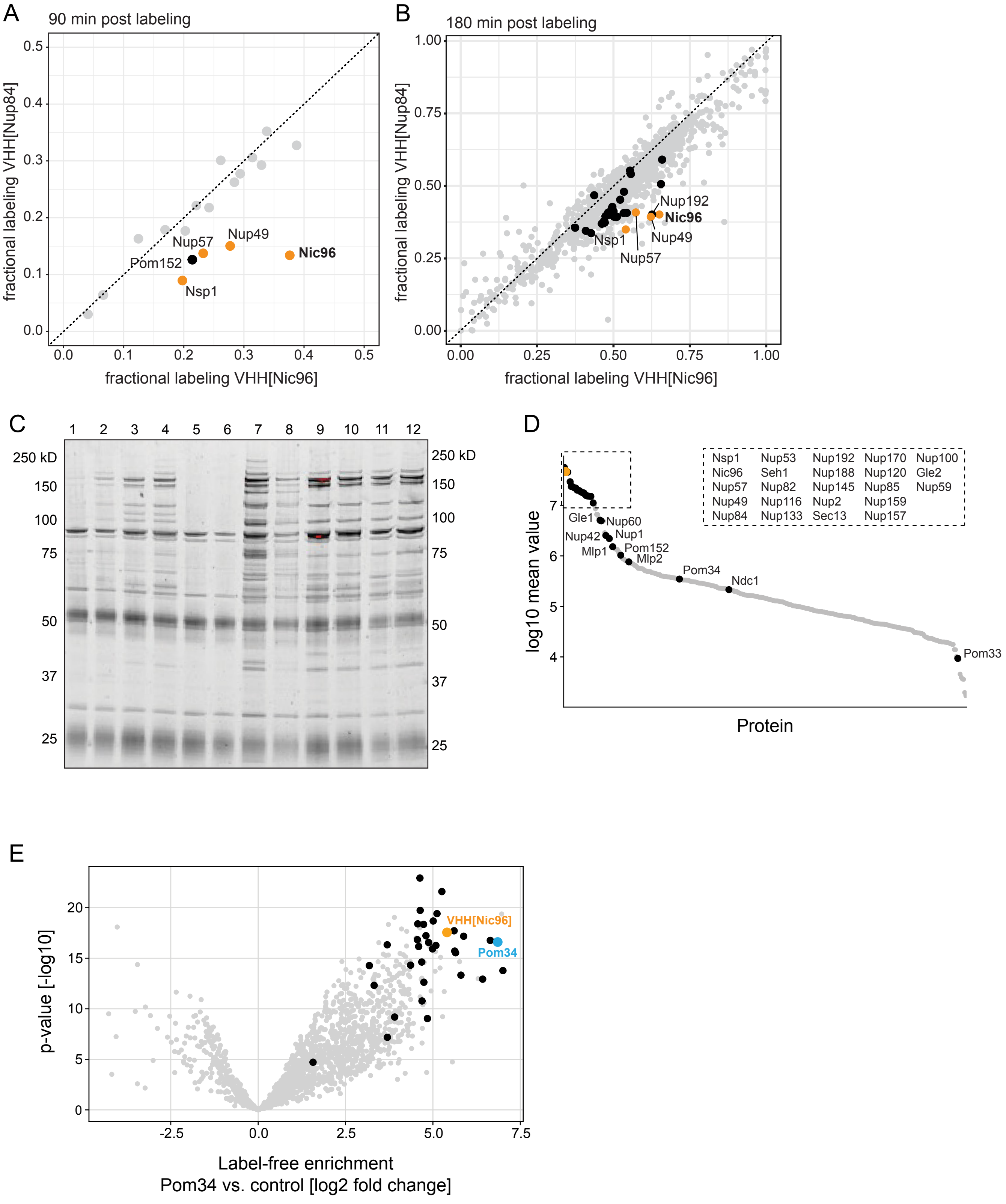

A
90 min post labeling
fractional labeling VHH[Nup84]
fractional labeling VHH[Nic96]
Nup57
Nup49
Nic96
Pom152
Nsp1
B
180 min post labeling
Nup192
Nic96
Nsp1
Nup49
Nup57
C
1 2 3 4 5 6 7 8 9 10 11 12
250 kD
150
100
75
50
37
25
D
Nsp1 Nup53 Nup192 Nup170 Nup100
Nic96 Seh1 Nup188 Nup120 Gle2
Nup57 Nup82 Nup145 Nup85 Nup59
Nup49 Nup116 Nup2 Nup159
Nup84 Nup133 Sec13 Nup157
Gle1
Nup60
Nup42
Nup1
Mlp1
Pom152
Mlp2
Pom34
Ndc1
Pom33
log10 mean value
Protein
E
VHH[Nic96]
Pom34
p-value [-log10]
Label-free enrichment
Pom34 vs. control [log2 fold change]


Figure S1

**Fig. S1 VHH[Nic96] binds newly synthesized Nic96 and direct Nic96 interaction partners**

**A,B**. Same data as in **Fig 1D,E**. Fractional labelling (Heavy / (Heavy + Light)) of all reproducibly co-purified proteins with VHH[Nup84] compared to VHH[Nic96] at t=90min (**A**) or t=180 (**B**). Each datapoint corresponds to the median fractional labelling of three biological replicates. Nups are coloured in black, the Nic96-containing CTN trimer is coloured in orange and all other proteins are plotted in grey.

**C.** Affinity purifications of constitutively expressed VHH[Nic96]-ZZ in various buffers to identify optimal conditions for co-purifying nups. Buffer compositions from [1]: 1) 20mM HEPES pH 7.4, 110mM K Acetate, 2mM MgCl2, 150mM NaCl, 1% Triton X-100, 0.1% Tween20, 2) buffer 1 + 10% glycerol, 3) 40mM Tris pH 8.0, 250mM $Na_3Cit$, 1% Brij 58, 300µM Sarkosyl, 4) buffer 3 + 10% glycerol, 5) 40mM Tris pH 8.0, 50mM $Na_3Cit$, 300mM NaCl, 0.1% Tween-20, 2mM EDTA, 6) buffer 5 + 10% glycerol, 7) 1.5M Ammonium Acetate pH 7.0, 1% Triton X-100, 8) buffer 7 + 10% glycerol, 9) 40mM Tris pH 8.0, 250mM $Na_3Cit$, 100mM NaCl, 1% Triton X-100, 10) buffer 9 + 10% glycerol, 11) 40mM Tris pH 8.0, 250mM $Na_3Cit$, 300mM NaCl, 1% Triton X-100, 12) buffer 11 + 10% glycerol. All buffers were additionally supplemented with 1mM DTT, PIC and 1:200 Solution P.

**D.** Nups co-purifying in VHH[Nic96]-ZZ affinity purifications shown in **Fig. 1F**. Each datapoint corresponds to mean fragment intensity identified from gel plugs of 3 replicates. Proteins are ranked in descending order. Nups are annotated in black; VHH[Nic96]-ZZ is shown in orange.

**E.** Nups and VHH[Nic96] co-purifying in Pom34-ZZ affinity purifications[2] compared to affinity purifications from cells expressing VHH[Nic96] but no ZZ-tag. Average enrichment of n=6 replicates. Nups are indicated in black, VHH[Nic96] is indicated in orange, Pom34 in blue. Pom34 and VHH[Nic96] are annotated.

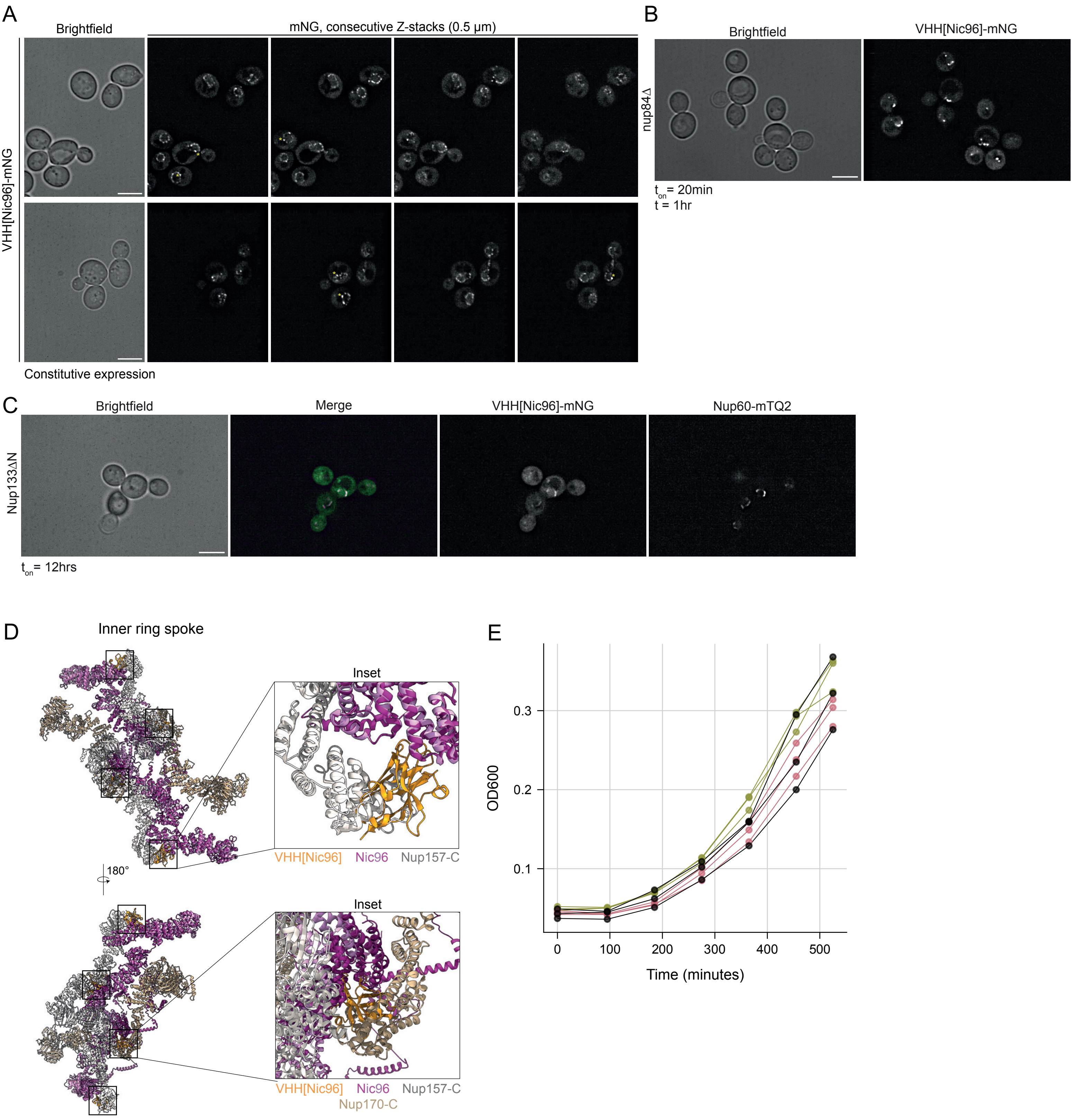
A
Brightfield
mNG, consecutive Z-stacks (0.5 μm)
VHH[Nic96]-mNG
Constitutive expression
B
Brightfield
VHH[Nic96]-mNG
nup84Δ
$t_{on}$ = 20min
t = 1hr
C
Brightfield
Merge
VHH[Nic96]-mNG
Nup60-mTQ2
Nup133ΔN
$t_{on}$ = 12hrs
D
Inner ring spoke
Inset
VHH[Nic96]
Nic96
Nup157-C
180°
Inset
VHH[Nic96]
Nic96
Nup157-C
Nup170-C
E
OD600
0.1
0.2
0.3
0
100
200
300
400
500
Time (minutes)

Figure S2

**Fig. S2 VHH[Nic96]-mNG is incorporated in NPCs during assembly**

**A.** Typical images of VHH[Nic96]-mNG constitutively expressed from a Nup120 promoter. Each panel represents a single z-slice in a consecutive series of z-slices (0.5 μm between slices). Brightness/contrast settings are identical to **Fig. 4A**. Asterisks indicate NVJ-proximal foci. Scale bar = 5 μm.

**B**. Localization of VHH[Nic96]-mNG in nup84Δ cells imaged at t=1hr following a 20-minute expression burst. Images represent single z-slices. Scale bar = 5 μm.

**C**. Localization of VHH[Nic96]-mNG in Nup133ΔN cells that co-express Nup60-mTurqoise2 (mTQ2). VHH[Nic96]-mNG was expressed overnight to allow visualization. Images represent single z-slices. Scale bar = 5 μm.

**D.** Model showing the position of VHH[Nic96] (orange) bound to Nic96 (purple / pink) in the inner ring spoke, showing the potential clash of VHH bound to Nic96 with the C-terminus of Nup157 (white) and the C-terminus of Nup170 (beige); based on the structure of Nic96 bound by VHH[Nic96] [3] and the inner spoke ring of the yeast NPC [4]. Please note VHH[Nic96] is not readily discernible due to its modelled localization within the inner ring spoke. Black squares mark the position of VHH[Nic96].

**E.** Growth curves of WT cells (green), VHH[Nic96]-mNG expressing cells (pink) and Nic96-mNG cells (black).

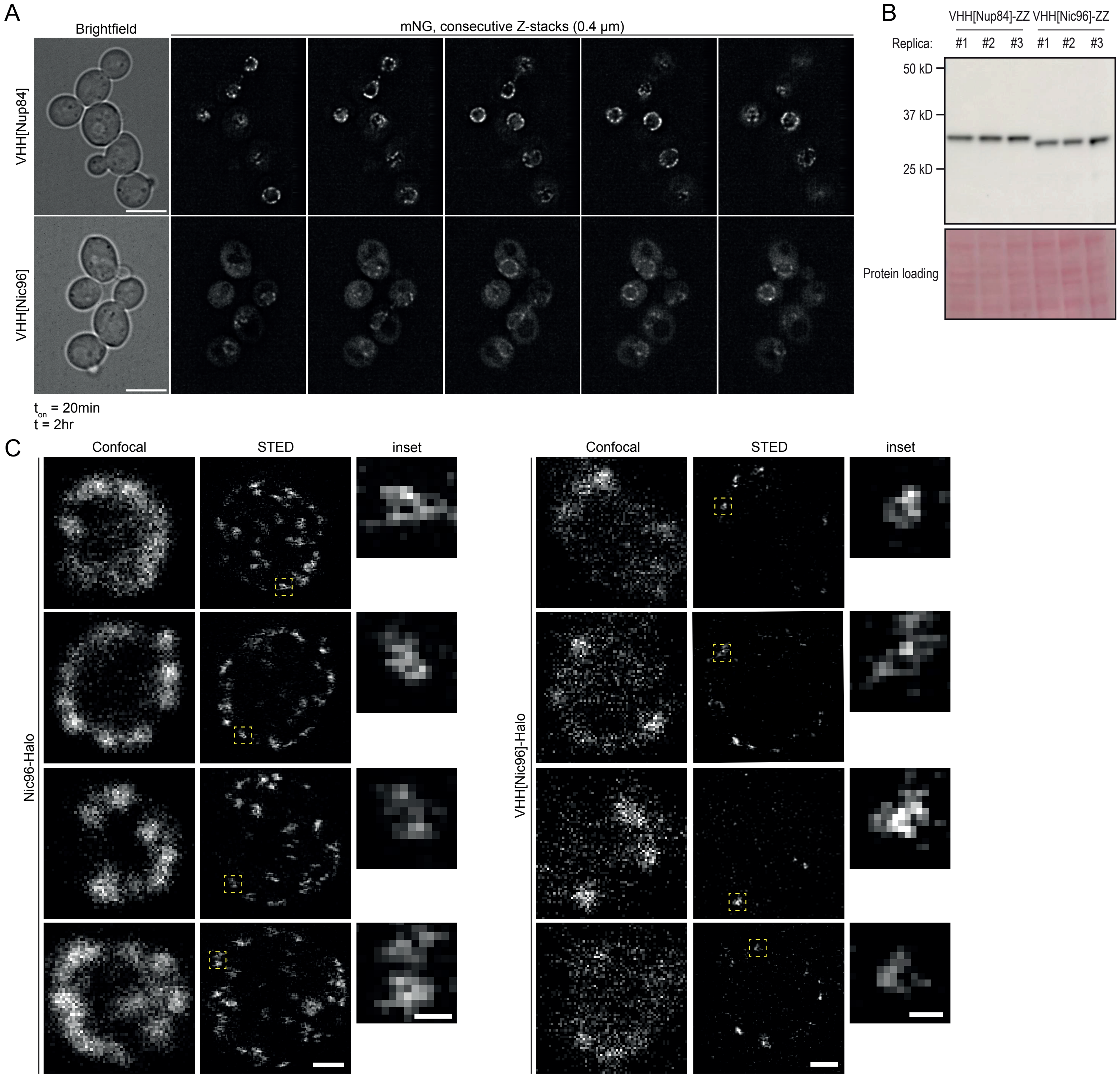
A
Brightfield
mNG, consecutive Z-stacks (0.4 µm)
VHH[Nup84]
VHH[Nic96]
t_on = 20min
t = 2hr
B
VHH[Nup84]-ZZ VHH[Nic96]-ZZ
Replica:
#1 #2 #3 #1 #2 #3
50 kD
37 kD
25 kD
Protein loading
C
Confocal
STED
inset
Nic96-Halo
VHH[Nic96]-Halo

Figure S3

**Fig. S3 Visualizing VHH[Nic96] shortly after its synthesis**

**A.** Typical images of the localization of both nanobodies at t=2hr following a 20-minute expression burst as described in **Fig. 2A**. Each panel represents a single z-slice in a consecutive series of z-slices (0.4 um between slices). Brightness/contrast settings are identical to **Fig. 4A and Fig. 5B**. Scale bar = 5 μm.

**B.** Representative immunoblot of cells expressing VHH-ZZ fusions after 90 minutes of expression. VHH[Nic96]-ZZ = 31.5 kD; VHH[Nup84]-ZZ = 33 kD. Data from three independent biological experiments.

**C.** Live-cell confocal and STED microscopy images of Nic96-Halo and VHH[Nic96]-Halo structures in the yeast nuclear envelope labelled with SiR-Halo dye. VHH[Nic96]-Halo expressing cells were imaged at t=1hr following a 20-minute expression burst. Insets indicate structures of approximately 100 nm. Scale bar = 0.1 μm (inset) and 0.5 μm (confocal).

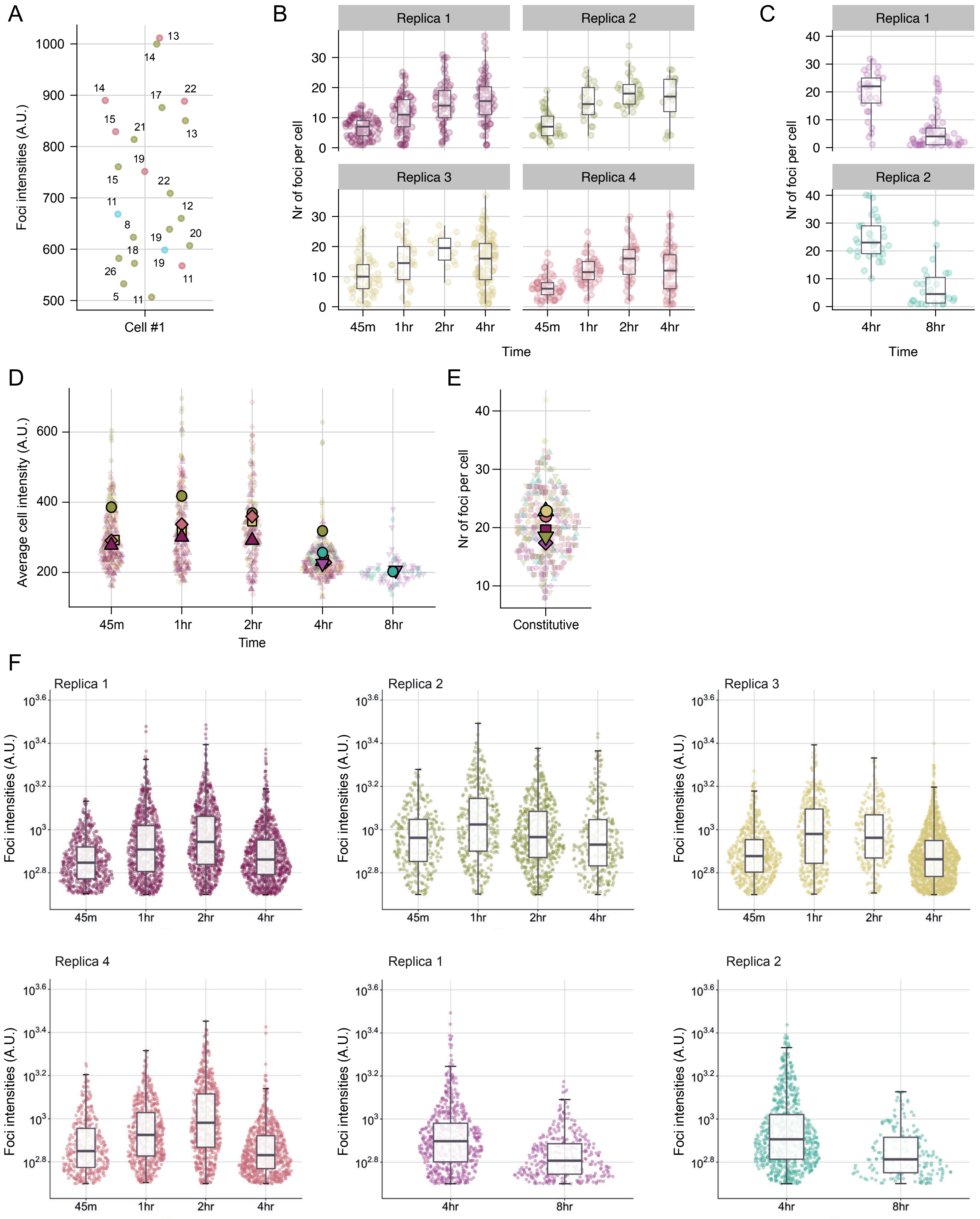
A
Foci intensities (A.U.)
Cell #1
B
Replica 1
Replica 2
Replica 3
Replica 4
Nr of foci per cell
Time
45m
1hr
2hr
4hr
C
Replica 1
Replica 2
Nr of foci per cell
4hr
8hr
Time
D
Average cell intensity (A.U.)
45m
1hr
2hr
4hr
8hr
Time
E
Nr of foci per cell
Constitutive
F
Replica 1
Replica 2
Replica 3
Replica 4
Foci intensities (A.U.)

Figure S4

**Fig. S4 Quantitative assessment of VHH[Nic96] labelling is consistent with dynamics of assembly and cell division - 1**

**A.** Foci intensities of detected VHH[Nic96]-mNG foci in the cell depicted in **Fig 5B**. Numbers indicate Z-slices, colours match squares in **Fig 5B**. A.U. = arbitrary units.

**B.** Number of foci per cell at each sampling timepoint in each replica. Each dot represents a cell, each replica reflects a single biological experiment. Overlayed boxplot depicts median and first and third quartiles. Whiskers extend to 1.5x IQR. Nr of cells analysed at t=45m, t=1hr, t=2hr and t=4hrs: *92, 75, 58, 64 (replica 1) - 39, 24, 31, 18 (replica 2) - 61, 30, 14, 107 (replica 3) - 58, 52, 44, 60 (replica 4)*.

**C.** Number of foci per cell at each sampling timepoint in each replica as in **B** but now at t=4hrs and t=8hrs. Nr of cells analysed at t=4hrs and t=8hrs: *31, 55 (replica 1) - 31, 26 (replica 2)*.

**D.** Number of foci when VHH[Nic96] is constitutively expressed from a Nup120 promoter. Each dot represents a cell, biological replicas are color-coded. Mean nr of foci per replica are overlayed.

**E.** Average cell intensities at indicated timepoints. Each dot represents a cell, colours reflect biological replicates, biological replicas are color-coded. Mean nr of foci per replica are overlayed.

**F.** Foci intensities of foci identified in cells from panels **B** and **C** at each sampling timepoint. Each dot represents a single focus. Overlayed boxplot depicts median and first and third quartiles. Whiskers extend to 1.5x IQR. A.U. = arbitrary units.

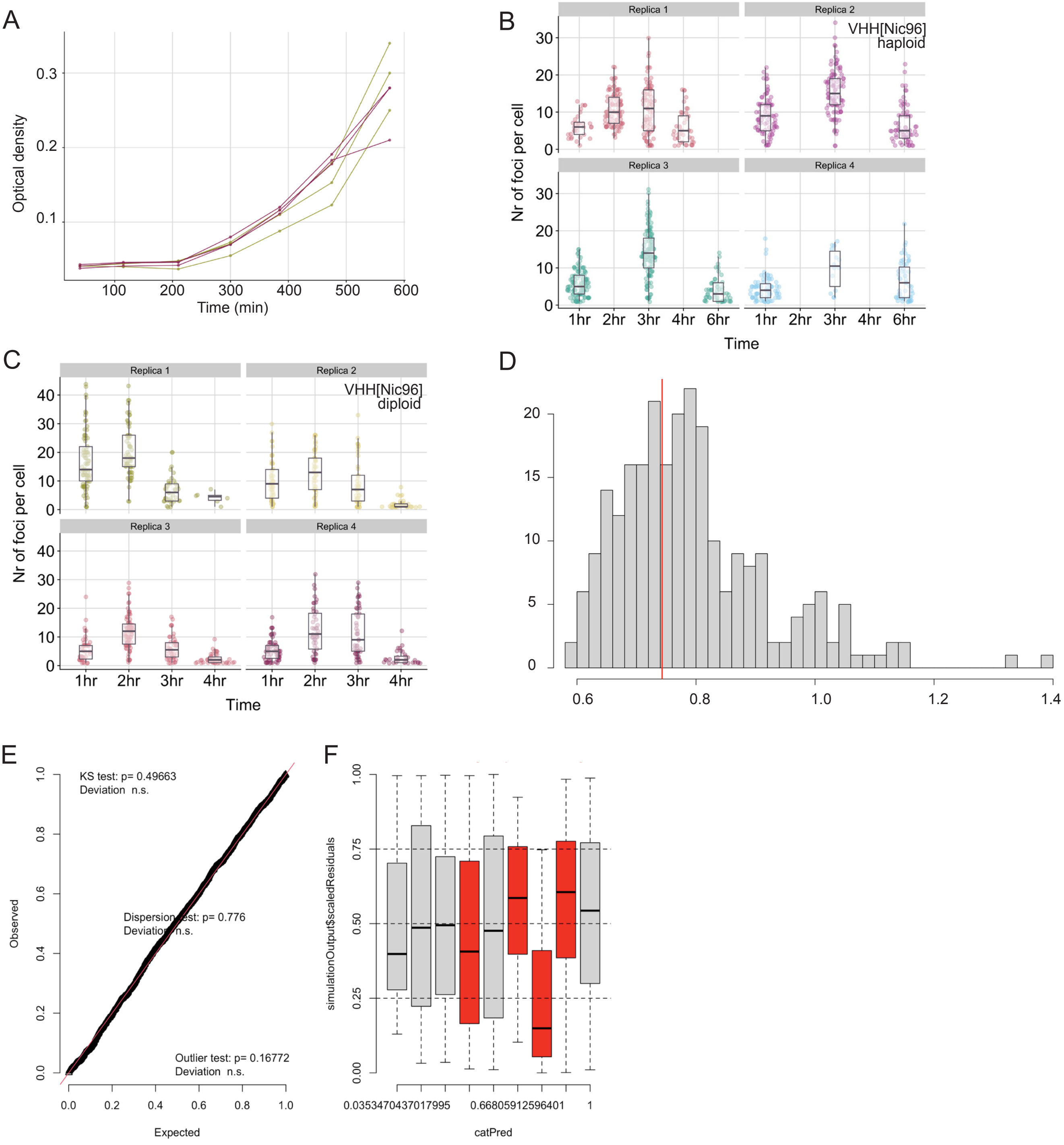
A
Optical density
Time (min)
B
Replica 1
Replica 2
Replica 3
Replica 4
VHH[Nic96]
haploid
Nr of foci per cell
1hr 2hr 3hr 4hr 6hr
Time
C
Replica 1
Replica 2
Replica 3
Replica 4
VHH[Nic96]
diploid
Nr of foci per cell
1hr 2hr 3hr 4hr
Time
D
E
KS test: p= 0.49663
Deviation n.s.
Dispersion test: p= 0.776
Deviation n.s.
Outlier test: p= 0.16772
Deviation n.s.
Observed
Expected
F
simulationOutput$scaledResiduals
0.0353470437017995
0.66805912596401
1
catPred

Figure S5

**Fig. S5 Quantitative assessment of VHH[Nic96] labelling is consistent with dynamics of assembly and cell division - 2**

**A.** Growth curves of haploid VHH[Nic96]-mNG expressing cells (green lines) and diploid VHH[Nic96]-mNG expressing cells (purple lines).

**B.** Number of foci per cell at each sampling timepoint in each replica in haploid strains. Each dot represents a cell; each replica reflects a single biological experiment. Overlayed boxplot depicts median and first and third quartiles. Whiskers extend to 1.5x IQR. Nr of cells analysed at t=1hr, t=2hr, t=3hr and t=4hrs: *77, 61, 40, 6 (replica 1) - 53, 49, 49, 30 (replica 2) - 42, 83, 44, 41 (replica 3) - 59, 44, 49, 32 (replica 4)*.

**C.** Number of foci per cell at each sampling timepoint in each replica in diploid strains. Each dot represents a cell; each replica reflects a single biological experiment. Overlayed boxplot depicts median and first and third quartiles. Whiskers extend to 1.5x IQR. Nr of cells analysed at t=1hr, t=2hr, t=3hr and t=4hrs: *32, 90, 82, 41 (replica 1).* Nr of cells analysed at t=1hr, t=3hr and t=4hrs: *81, 93, 65 (replica 2) - 111, 123, 53 (replica 3) - 74, 14, 56 (replica 4).*

**D.** DHARMa diagnostic plot of the scaled residuals for the fitted models depicted in **Fig. 5F**. Simulated values, red line = fitted model. P-value (two sides) = 0,776.

**EF.** DHARMa residuals. QQ plot residuals (left); Within-group deviation from uniformity significant (red), Levene Test for homogeneity of variance significant (right).

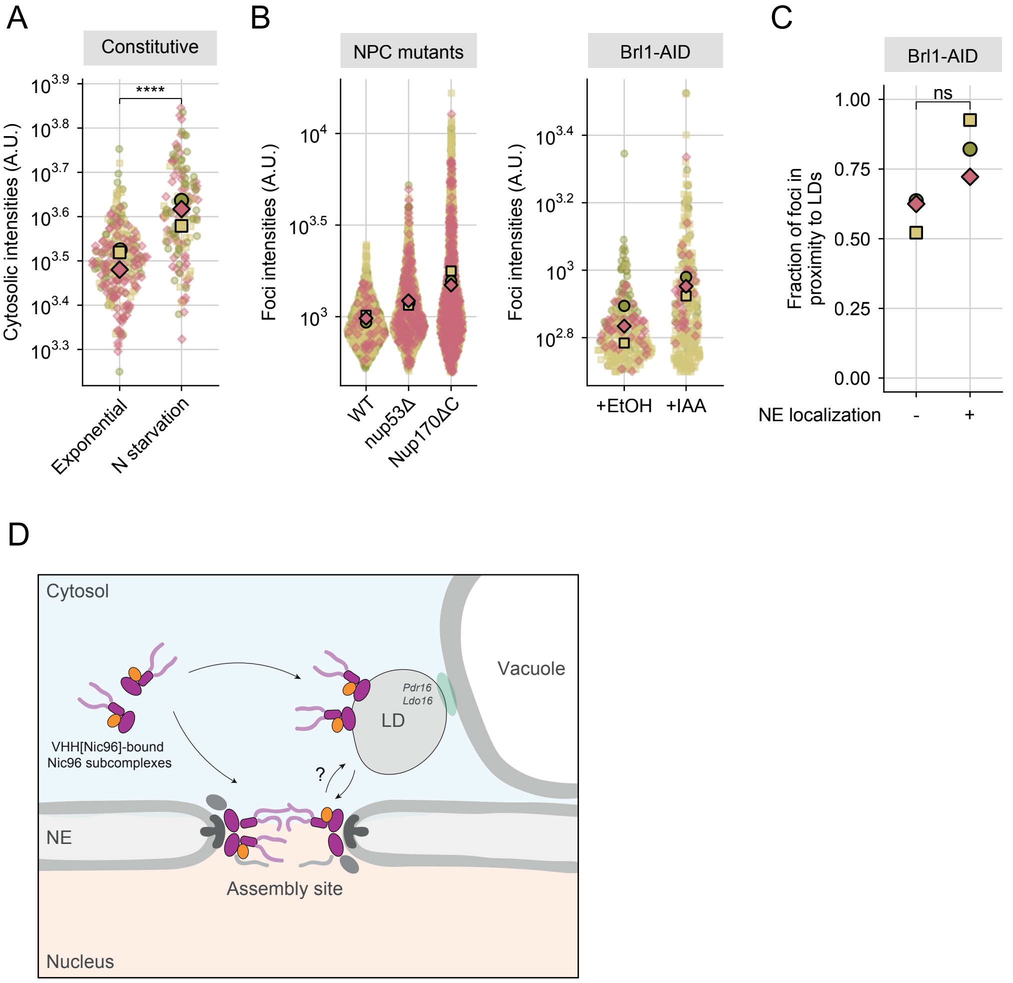

A
Constitutive
Cytosolic intensities (A.U.)
10^3.9
10^3.8
10^3.7
10^3.6
10^3.5
10^3.4
10^3.3
****
Exponential
N starvation
B
NPC mutants
Foci intensities (A.U.)
10^4
10^3.5
10^3
WT
nup53Δ
Nup170ΔC
Brl1-AID
Foci intensities (A.U.)
10^3.4
10^3.2
10^3
10^2.8
+EtOH
+IAA
C
Brl1-AID
ns
1.00
0.75
0.50
0.25
0.00
Fraction of foci in proximity to LDs
NE localization
-
+
D
Cytosol
Vacuole
Pdr16
Ldo16
LD
VHH[Nic96]-bound Nic96 subcomplexes
?
NE
Assembly site
Nucleus


Figure S6

**Fig. S6 Association of Nic96 subcomplexes with NE-localized lipid droplets**

**A.** Quantification of cytosolic VHH[Nic96]-mNG intensities in exponentially growing cells and cells starved for nitrogen for a duration of 2hrs. Cells constitutively express VHH[Nic96]-mNG from a Nup120 promoter. Each dot represents a cell, color reflects biological replica with total nr of cells > 130 per condition. Means of each replicate are overlayed. **** $p < 0.001$; *Welch's T-test on log-transformed data.*

**B. Left:** VHH[Nic96]-mNG foci intensities at t=2hr following a short, 20-minute induction pulse in WT, nup53Δ and Nup170ΔC cells. **Right:** VHH[Nic96]-mNG foci intensities at t=1hr following a short, 20-minute induction pulse in control (EtOH) and Brl1 depleted (Brl1-AID) cells. Brl1 was depleted as in Fig. 2E. Replica means are overlayed and biological replica are color-coded. Each dot represents a single focus. A.U. = arbitrary units. Nr of foci analysed: *441, 254, 53 (WT) - 372, 506, 225 (nup53Δ) – 493, 1118, 485 (Nup170ΔC) - 91, 152, 42 (EtOH) - 55, 211, 33 (Brl1-AID)*

**C.** Quantification of LD proximity of VHH[Nic96]-mNG foci after Brl1 depletion. Same data as in **Fig. 6I** but now stratified by NE localization. Biological replica are color-coded. Statistical significance was assessed using a binomial generalized linear model with predictor effects evaluated using likelihood ratio tests (ANOVA). $p < 0.1$ (ns); $p = 0.07$.

**D.** Working model for the balance between NPC assembly and storage on NE-localized lipid droplets that is supported by the data in this manuscript. VHH[Nic96]-bound Nic96 subcomplexes are incorporated into new NPCs but can also be deposited on lipid droplets (LD) close to the NE. The association with lipid droplets is favoured when assembly is perturbed. A subpopulation of these LDs is Pdr16- and Ldo16-positive and resides near the NVJ [5,6], establishing a contact site with the vacuole. In unperturbed conditions, we observe proximity between NE-localized VHH and LDs, which may reflect a close proximity of the assembly site to LD sites.

**Supplementary Table 1:**

Analysis of Deviance Table (Type II Wald chisquare tests) for model 1 that describes the number of VHH[Nic96] foci over time (Fig. 5F). [1)]

**Model 1: no_foci ~ strain_no* time + strain_no*I(time^2) + (1 | replica)**

**Response: no_foci**

| | **Chisq** | **Df** | **Pr(>Chisq)** |
|---|---|---|---|
| strain_no | 0.0484 | 1 | 0.8258 |
| time_c | 334.5073 | 1 | < 2.2e-16 *** |
| I(time_c^2) | 346.1814 | 1 | < 2.2e-16 *** |
| strain_no:time_c | 29.7893 | 1 | 4.816e-08 *** |
| Strain_no:I(time_c^2) | 98.7863 | 1 | < 2.2e-16 *** |

[1)] To evaluate the relationship between the number of foci in a cell and time post induction pulse in diploid and haploid cells, we built a range of candidate models and selected the final model based on the best AIC. No_foci = number of foci.

Strain_no = strain background (haploid / diploid). Replica = unique identifier for each combination of date and strain_no. Time = total time after induction pulse following the imaging set-up described in **Fig. 2A**.

**Supplementary Table 2:**

Estimated coefficients for the model from Supplementary table 1.

| | Coefficient | Estimate | Standard Error |
|---|---|---|---|
| intercept | $\beta_{\mathrm{intercept}}$ | 1.144057 | 0.127014 |
| Effect of strain (diploid vs. haploid) | $\beta_{\mathrm{diploid}}$ | -0.11276 | 0.207007 |
| Linear effect of time (haploid) | $\beta_{\mathrm{t}}$ | 0.842886 | 0.05922 |
| Quadratic effect of time (haploid) | $\beta_{\mathrm{t2}}$ | -0.1251 | 0.008496 |
| Strain x time interaction | $\beta_{\mathrm{t:diploid}}$ | 0.759697 | 0.139191 |
| Strain x time$^2$ interaction | $\beta_{\mathrm{t2:diploid}}$ | -0.27882 | 0.028053 |
| Dispersion parameter | | 3.16 | |
| Sd random effect (replica) | | 0.1939 | |

1 **Hakhverdyan, Z. *et al.* Rapid, optimized interactomic screening. *Nature Methods* 12, 553–560 (2015). https://doi.org/10.1038/nmeth.3395**

2 **Alber, F. *et al.* The molecular architecture of the nuclear pore complex. *Nature* 450, 695–701 (2007). https://doi.org/10.1038/nature06405**

3 **Nordeen, S. A. *et al.* A nanobody suite for yeast scaffold nucleoporins provides details of the nuclear pore complex structure. *Nat Commun* 11, 6179 (2020). https://doi.org/10.1038/s41467-020-19884-6**

4 **Akey, C. W. *et al.* Implications of a multiscale structure of the yeast nuclear pore complex. *Molecular Cell* 83, 3283–3302.e3285 (2023). https://doi.org/10.1016/j.molcel.2023.08.025**

5 **Diep, D. T. V. *et al.* A metabolically controlled contact site between vacuoles and lipid droplets in yeast. *Developmental Cell* 59, 740–758.e710 (2024). https://doi.org/10.1016/j.devcel.2024.01.016**

6 **Álvarez-Guerra, I. *et al.* LDO proteins and Vac8 form a vacuole-lipid droplet contact site to enable starvation-induced lipophagy in yeast. *Developmental Cell* 59, 759–775.e755 (2024). https://doi.org/10.1016/j.devcel.2024.01.014**
